# Gut microbial interaction networks control autoimmunity to neuroretina

**DOI:** 10.64898/2025.12.04.691931

**Authors:** Amy Zhang, Reiko Horai, Yingyos Jittayasothorn, Jonathan H. Badger, Zhichao Wu, Guangpu Shi, Akriti Gupta, Samyuktha Arunkumar, Caitlin E. Murphy, Vijayaraj Nagarajan, John A. McCulloch, Shilpa Kodati, H Nida Sen, Jung Wha Lee, Jonathan P. Jacobs, Xiaoyan Xu, Mary J. Mattapallil, Zixuan Peng, Biying Xu, Robert J. Palmer, Nadim Majdalani, Kenya Honda, Colm O’hUigin, Rachel R. Caspi

**Author notes:** Correspondence to: Rachel R. Caspi, Laboratory of Immunology, National Eye Institute, National Institutes of Health 10 Center Drive, 10/10N248, Bethesda, MD 20892-1857, USA, Reiko Horai, Laboratory of Immunology, National Eye Institute, National Institutes of Health 10 Center Drive, 10/10S243, Bethesda, MD 20892-1857, USA. Equal contribution (Amy Zhang, Reiko Horai).

## Abstract

The gut microbiome influences the development of immune-mediated inflammatory diseases, including autoimmune uveitis, a sight-threatening ocular inflammation driven by retina-specific T cells^1^. Using a model of spontaneous autoimmune uveitis (sEAU) we showed that gut commensals provide immune stimuli that trigger disease^2^. Here we report that uveitis-promoting microbes are present in human gut flora and that colonization of germ-free (GF) mice with commensals from healthy human donors was sufficient to provoke disease. Severity of sEAU correlated with expansion of Akkermansia and contraction of short-chain fatty acid (SCFA)-producing Firmicutes, accompanied by decreased SCFA levels and a dominant gut Th1 effector response. Mechanistic gain-of-function experiments, enriching GF sEAU mice with *A. muciniphila*, reproduced these microbiome, metabolite and immune phenotype shifts, and exacerbated disease. We propose that Akkermansia promotes autoimmunity by outcompeting SCFA producers and enhancing Th1-type responses. An inverse correlation between Akkermansia (Verrucomicrobia) and Firmicutes was also present in fecal microbiota from patients with uveitis; related Verrucomicrobia/Firmicutes shifts were observed in multiple sclerosis and Crohn’s disease datasets. These findings reveal a stereotypic gut microbial interaction network that regulates systemic immune balance, and may represent an ecologically conserved mechanism through which the gut microbiome modulates autoimmune and inflammatory diseases.

## INTRODUCTION

Immune-privileged tissues are increasingly recognized as sites where host–microbiota interactions influence immunological balance in autoimmune and inflammatory diseases^3,4^. Multiple mechanisms have been proposed that explain the central role of microbiota in driving inflammation in the central nervous system (CNS), including innate and adaptive effects such as molecular mimicry^2,5^, bystander activation^6^, epitope spreading^7,8^ and dual T cell receptors (TCRs)^9^. The eye, especially the neuroretina, is part of the CNS and represents a quintessential immune-privileged tissue, making it a valuable model to study immune privilege and autoimmunity in the CNS.

Autoimmune uveitis is a T cell-driven intraocular inflammation of unknown etiology that targets the neuroretina and can lead to blindness^1^. It is represented by the models of experimental autoimmune uveitis (EAU) in animals. EAU is induced in wild type mice by immunization with the retinal antigen IRBP^1^, or can develop spontaneously (sEAU) in mice expressing a transgenic IRBP-specific TCR cloned from the immunization induced model (line R161H)^10^. Our previous studies highlighted the critical role of gut microbiome in triggering sEAU by providing antigenic stimuli that activate retina-specific T cells^2^.

Subsequent research reported inhibitory effects of microbiome depletion also in induced EAU^11^, where the antigen is provided, emphasizing the antigen-independent (innate?) effects of gut microbes on EAU. Interestingly, gut microbiome studies in uveitis patients have revealed differences in gut flora between patients and healthy controls^12–16^, but mechanistic information could not be resolved.

Because inductive events and causal relationships cannot be studied in patients, we employed the model of sEAU in R161H TCR transgenic mice^10^, which may provide a more physiological system to study triggers delivered by the microbiota. To gain mechanistic insights into how human gut microbiota might affect the development of autoimmune uveitis, we created gnotobiotic breeding colonies of R161H mice harboring healthy human gut flora by reconstituting GF parental mice with fecal material from three healthy human donors. We focused on healthy donors because individuals who develop uveitis are clinically healthy before disease onset; disease-promoting organisms must therefore already be present during this period. Microbiota obtained after uveitis develops may be altered by disease or treatment, confounding interpretation of initiating events.

Using a multiomic strategy that integrates microbial, metabolomic and immune profiling of human flora (HuFl)-reconstituted R161H mice, combined with analyses of human autoimmune disease datasets, we identified a microbial interaction network comprising an “Akkermansia (Verrucomicrobia)—Firmicutes axis” that appears to regulate autoimmunity. Our data points to an underexplored but fundamental mechanism, where interactive relationships in the microbial community play a central role in regulation of autoimmune disease pathogenesis by microbiota^17^.

## RESULTS

### 1. Human gut commensals support development of spontaneous experimental autoimmune uveitis (sEAU)

R161H mice harboring mouse microbiota (specific-pathogen-free, SPF) typically develop moderate to severe uveitis with high penetrance (100% incidence by two months of age)^10^. To first assess whether human gut microbiota affect uveitis development, we developed “humanized gnotobiotic” R161H mouse lines as our pre-clinical model, by reconstituting breeding pairs of GF R161H mice with gut microbiota from three healthy human subjects—U, V, and W. Each line was maintained in separate isolators through successive generations, so that the offspring acquired the human flora naturally from their respective parents (Fig. 1a).

**Fig. 1.**
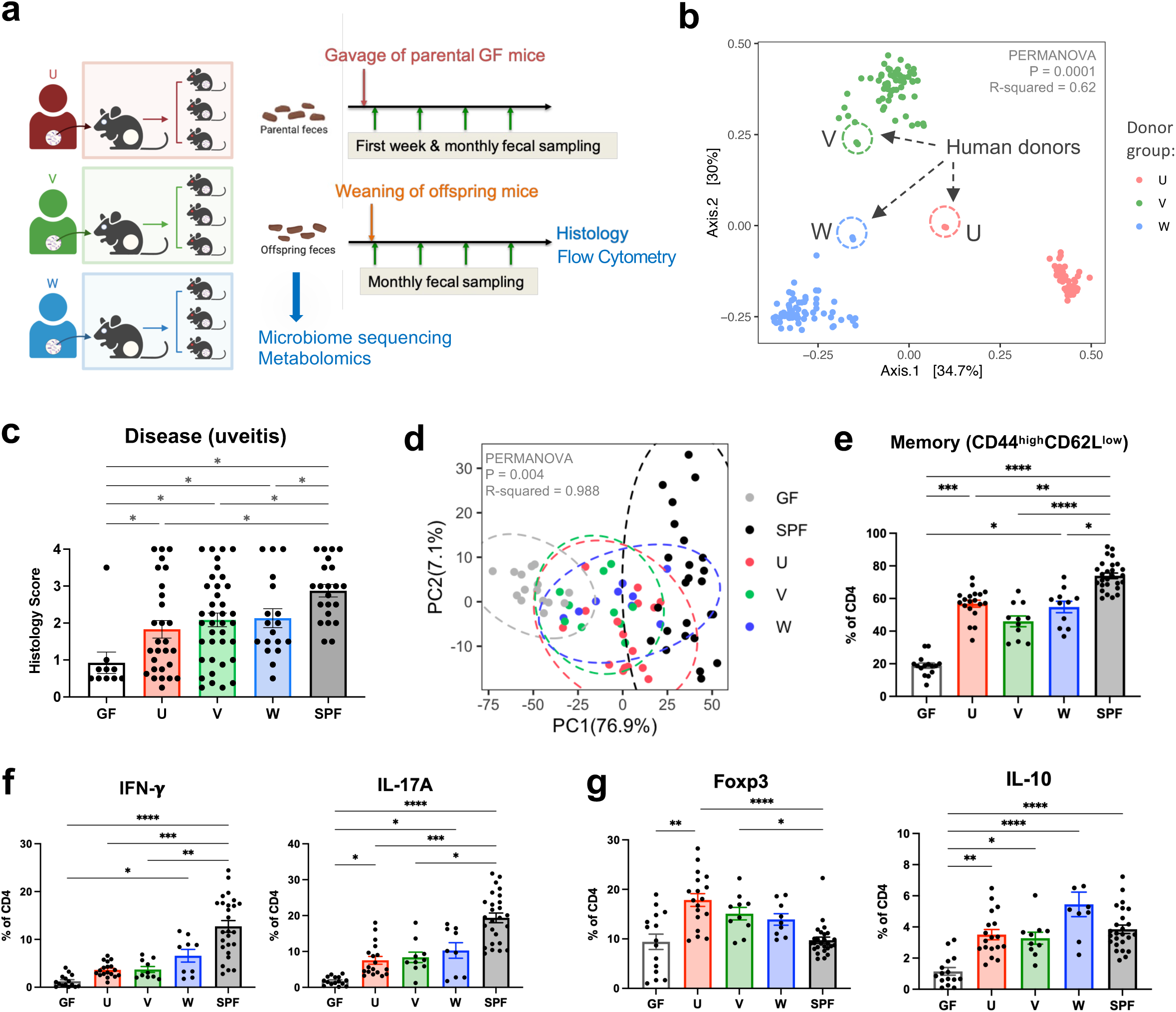
Human gut commensals support development of spontaneous experimental autoimmune uveitis (sEAU). **a**, Schematic representation of human fecal microbiota transfer experiments and data collection. **b**, Beta diversity based on Bray–Curtis dissimilarity showing differences in fecal microbiome composition between recipient groups U (*n =* 50), V (*n =* 59), W (*n =* 64). Encircled with dashed lines are the respective human donor (original) fecal samples. **c**, Disease scores (histology) of age-matched R161H mice (4-5 months old) from human flora (U, V, W, *n =* 28, 38, 18), SPF (*n =* 22) and GF (*n =* 10) groups (Kruskal–Wallis test between mouse recipient groups, p = 0.0002). **d**, Immunological clusters of CD4 T cells from colon lamina propria (LP) by the microbiome groups. PCA analysis performed on immune signature matrix by frequencies of CD4 T cells expressing each cell surface or intracellular marker (see **Extended Data Fig. 1e**). **e**–**g**, Frequencies of memory (CD44^high^CD62L^low^, **e**), IFN-*γ*-or IL-17A-producing (**f**), and regulatory (Foxp3-positive or IL-10-producing, **g**) CD4 T cells in colon LP of HuFl, SPF and GF R161H mice. **d**–**g**, Compiled data from 12 experiments (2-3 months old), *n =* 15, 19, 11, 11, 27 for GF, U, V, W and SPF, respectively. Kruskal–Wallis tests show p < 0.0001 (**e**–**g**). P-values adjusted for post hoc multiple comparisons are displayed as asterisks. *P < 0.05, **P < 0.01, ***P < 0.001, ****P < 0.0001, and bar plots showing mean values ± SEMs (**c**, **e**–**g**).

Microbiome profiling confirmed the establishment of human donor-derived microbial communities in gnotobiotic R161H mice. Although cross-species transfer resulted in selective engraftment, recipient communities remained distinguishable according to donor origin across successive generations in gnotobiotic isolators. The three cohorts thus retained donor-associated differences in community structure despite adaptation to the murine host (Fig. 1b).

Human commensal microbes supported uveitis development in R161H mice. Although HuFl mice display a broad spectrum of disease scores, the scores were on average lower than those of SPF mice that harbor normal mouse flora (Fig. 1c & Extended Data Fig. 1a). We did not observe obvious sex differences with respect to disease severity in HuFl mice (Extended Data Fig. 1b). Immune phenotype of colonic CD4^+^ T cells, based on a range of cell surface and intracellular markers, clearly distinguished HuFl mice from GF and SPF mice (Fig 1d & Extended Data Fig. 1c-e). The HuFl mice showed a spectrum of disease scores and memory marker expression between GF (low) and SPF (high) mice (Fig. 1e), had fewer IFN-*γ*-producing Th1 and IL-17A-producing Th17 effector cells and more regulatory T cells (Tregs) expressing Foxp3 and IL-10 than did SPF mice (Fig. 1f-g). This immune phenotype was consistent with other activation markers and Th1/Th17 signature molecules (Extended Data Fig. 1e). This phenotype of less effector and more regulatory cells present in the gut may explain the lower disease scores that they developed.

Because the central aim of this study was to define how human-derived gut microbial communities influence spontaneous autoimmunity, subsequent microbiome–immune association analyses focused on the HuFl cohorts. SPF mice served as a conventional reference for disease and immune phenotypes but were not included in these analyses, as their endogenous murine microbiota represents a distinct ecological context from the human donor-derived communities.

### 2. HuFl sEAU tracks with a dominant Th1 phenotype and abundance of Verrucomicrobia/Akkermansia

Since both Th1 and Th17 effector phenotypes can drive autoimmune uveitis^18–20^, we set out to define the immune signatures that associate with disease in the HuFl model. We categorized histological scores into high (score ≥ 1.5), mid (0.75 < score < 1.5), and low (score ≤ 0.75) to facilitate statistical analysis. Interestingly, IFN-*γ*-producing CD4 T cells, rather than IL-17A producers, in the gut (and other tissues) associated strongly with severe disease phenotype (Fig. 2a-d). More severe uveitis in HuFl mice was associated with a Th1-like response profile.

**Fig. 2.**
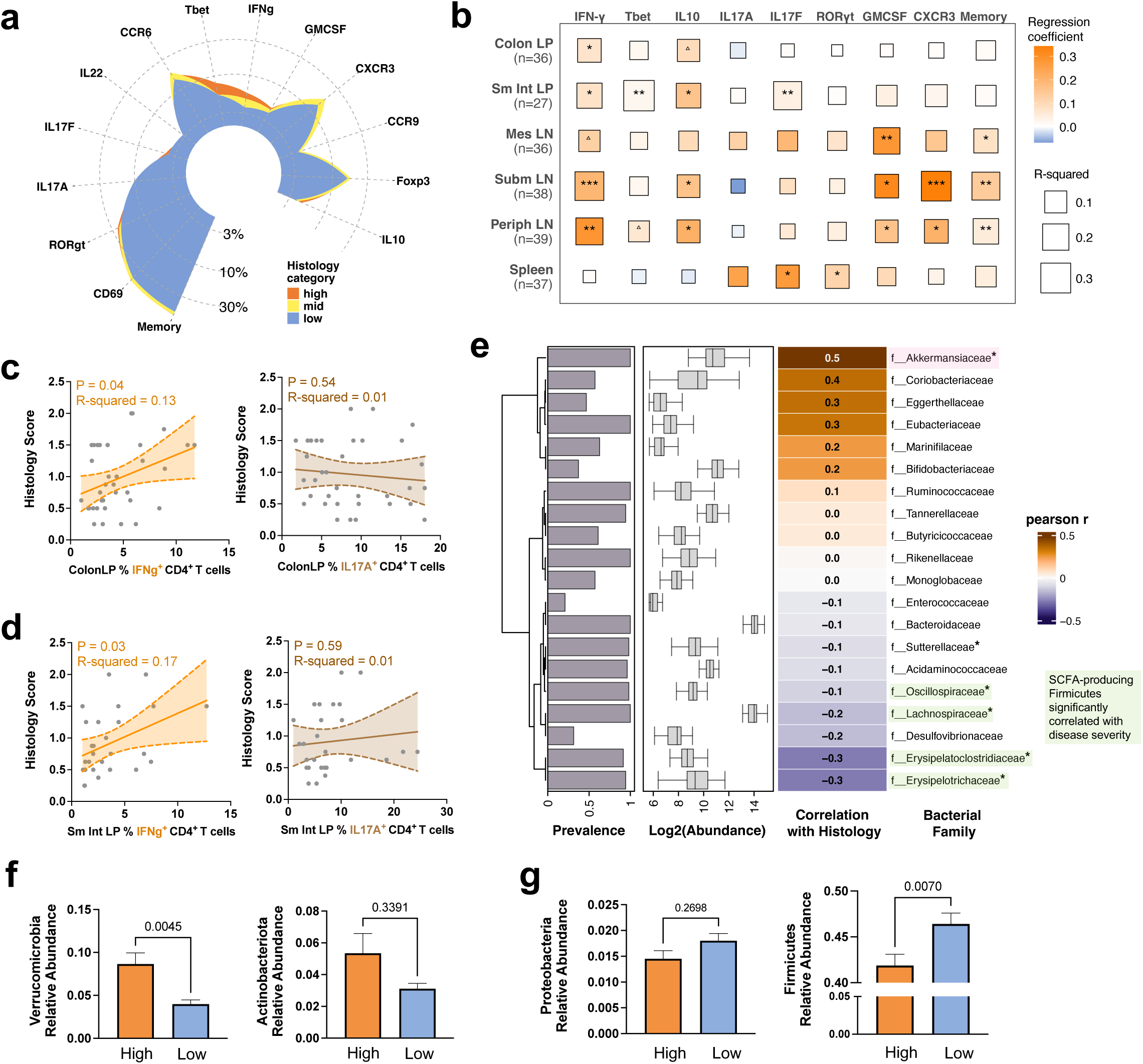
Human flora (HuFl) sEAU tracks with a dominant Th1 phenotype and abundance of Akkermansia (Verrucomicrobia). **a**, Mean relative frequency of colon LP CD4 T cells of each cell surface or intracellular marker arranged in a circular plot, colored by histology category: high (score ≥ 1.5, *n =* 10), mid (0.75 < score < 1.5, *n =* 8), low (score ≤ 0.75, *n =* 17). **b**, Correlational heatmap of histology scores with relative frequencies of each cell surface or intracellular marker in CD4 T cells across six tissue sites. LP, lamina propria; LN, lymph nodes; Sm Int, small intestine; Mes, mesenteric; Subm, submandibular; Periph, peripheral. ^△^0.05 < P < 0.08, *P < 0.05, **P < 0.01, ***P < 0.001, modeled by simple linear regression. **c**–**d**, Representative trend lines with 95% confidence interval (shaded area) show that disease scores of HuFl mice are positively associated with IFN-*γ*-producing, but not IL-17A-producing, CD4 T cells, in colon LP (**c**) and small intestine LP (**d**). Data in **a**–**d** are aggregated from 8 experiments (UVW combined, sample sizes ranged from *n =* 27 to 39 across tissues following quality filtering). **e**, Correlational heatmap showing major fecal bacterial taxa (out of top 20 most abundant families) that correlate with disease scores in adult HuFl R161H (UVW combined, *n =* 105). P-values assessed after FDR (false discovery rate) correction following Pearson correlation. *P < 0.05. The highlighted SCFA-producing Firmicutes are the families among the 20 most abundant that met the significance threshold. **f**–**g**, More Verrucomicrobia (**f**), and fewer Firmicutes (**g**) are present in the gut of HuFl mice with high disease scores. Sample sizes are *n* = 36 and 43 for high and low histology categories, respectively. Data in bar plots are presented as mean values ± SEMs, and P-values from Mann-Whitney U tests are shown in plots.

Immune phenotyping focused on the gut lamina propria and lymphoid tissues, where microbiota-associated immune priming occurs. Eyes were reserved for histopathological assessment and were therefore unavailable for flow-cytometric analysis of infiltrating T cells.

Microbiome analysis of parental and offspring HuFl mice showed that colonized mice adopted only a subset of the original donor taxa (Extended Data Fig. 2a-b). This could be a result from loss during human stool sample preservation and transfer processes, and/or from inefficient engraftment of some non-native microbes to the gut environment of a different host species. However, in terms of the overall microbial composition, each mouse cohort remained closer to their original donor, preserving a distinct microbial footprint (Fig. 1b & Extended Data Fig. 2c-d). Differences between female *vs.* male HuFl mice in terms of microbial abundance and composition were minimal, if any, compared to the differences based on human donors (Extended Data Fig. 2e-f). This is compatible with the apparent lack of sex differences in disease severity of HuFl mice (Extended Data Fig. 1b).

We then explored how gut microbiome associates with severity of uveitis. Correlational heat map of major bacterial taxa at the family level revealed strong links between several taxa and histology scores (Fig. 2e). Among them, Akkermansia (the only representative of Verrucomicrobia in HuFl mice) correlated with high disease scores (Pearson r = 0.5, FDR <0.05), whereas certain members of the Firmicutes phylum were correlated with low disease scores (Pearson r =-0.3, FDR <0.05). Differential abundance analysis between high and low histology categories consistently showed that Akkermansia was enriched in mice with high disease, whereas Firmicutes (especially Clostridia) were enriched in mice with low disease (Fig. 2f-g & Extended Data Fig. 2g).

### 3. SCFA signatures are enriched in low disease HuFl mice or healthy subjects

SCFAs are metabolites of commensal gut microbes resulting from fermentation of dietary fiber, and have anti-inflammatory properties in a range of pathological conditions^21^. Some SCFAs, e.g. butyrate and propionate, may have therapeutic potential in induced EAU^22,23^. In the current study, a key observation from differential abundance analyses of mice with low and high disease scores was that SCFA-producing Firmicutes were associated with lower sEAU scores (Fig. 2e). Therefore, it appeared plausible that SCFAs may also have the potential to modulate sEAU.

To address this, we mined functional gene signatures from shotgun metagenomic sequences of HuFl mice. Genes known to be involved in SCFAs synthesis and metabolism were mapped to the simplified pathways shown in Fig. 3a. Consistent with the observed differences in microbial abundance (Fig. 2e), the genes involved in main SCFA pathways were enriched in mice with low disease scores and reduced in mice with high disease scores (Fig. 3b). In agreement with the pathway analysis, metabolomic quantification of SCFAs in the gut and serum of the mice confirmed that higher disease scores were associated with lower SCFA concentrations both locally and systemically (Fig. 3c).

**Fig. 3.**
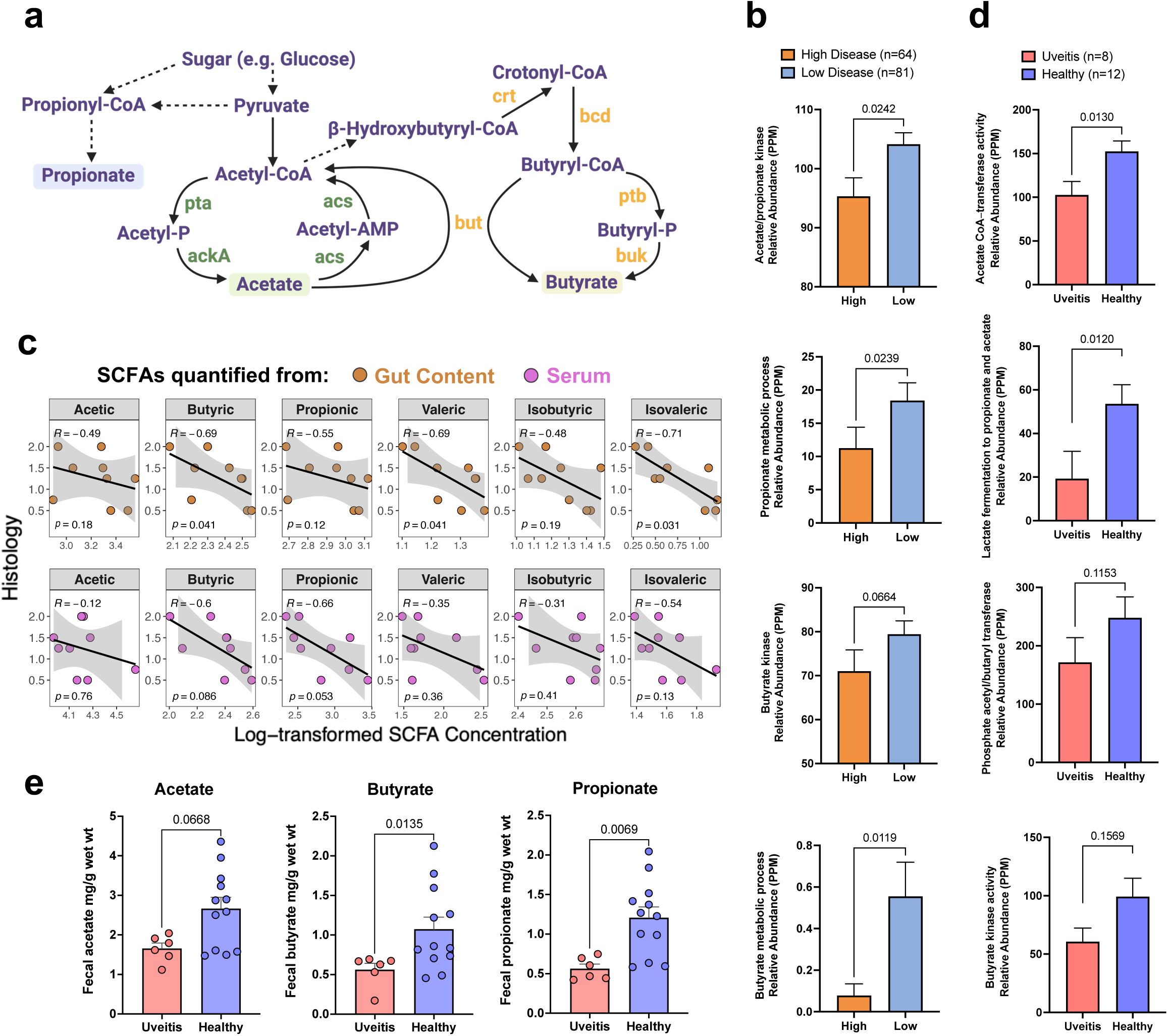
SCFA signatures are enriched in low disease HuFl mice or healthy subjects. **a**, A simplified pathway map highlighting genes associated with the major differentially abundant SCFA synthesis and metabolic processes; the schematic is not intended to represent all known SCFA-producing pathways. Dashed arrows indicate hidden intermediate steps. Abbreviations of enzyme symbols: pta, Phosphotransacetylase; ackA, Acetate kinase; acs, Acetyl-CoA synthetase; but, Butyryl-CoA:acetate CoA-transferase; ptb, Phosphate butyryltransferase; buk, Butyrate kinase; crt, Crotonase; bcd, Butyryl-CoA dehydrogenase. **b**, Enzymes in SCFA pathways (annotated with InterPro and Gene Ontology databases) are enriched in HuFl mouse gut with low histology scores (UVW combined, weanling and adult samples included). PPM, parts per million, a measure of relative abundance based on normalized base count for each feature in a metagenomic sample. **c**, Spearman correlation (visualized with a linear trend line) showing higher disease scores associated with lower SCFA concentrations both locally (in gut content) and systemically (in serum) in representative cohort U mice (*n =* 9). **d**, Enzymes in SCFA pathways (annotated with Gene Ontology and Pfam databases) are enriched in fecal samples from healthy humans compared to those from treatment-naïve uveitis patients, from a cohort of autoimmune uveitis (ocular-restricted, including idiopathic uveitis, VKH/SO and BCR). Sex, age, race and diet are comparably represented in patient (*n =* 8) and control (*n =* 12) groups. Normalized gene abundance is reported in PPM. **e**, SCFA levels from uveitis patients and healthy controls (same cohort as in **d,** but *n =* 6 available from the patient group). For **b, d** & **e**, Data are presented as mean values ± SEMs. Statistical significance was determined by Mann-Whitney U test.

We next examined whether this signature was present in a cohort of uveitis patients seen at the National Eye Institute (NEI) clinic, compared to healthy controls. Because uveitis is a heterogenous disease, we focused on a subset of patients whose disease was restricted to the eye (idiopathic uveitis, Vogt-Koyanagi-Harada disease/sympathetic ophthalmia and birdshot chorioretinopathy), as modeled by EAU, rather than being part of a systemic syndrome. Similarly to the HuFl R161H mice, enzymes involved in acetate, butyrate and propionate pathways (Fig. 3d) and their corresponding SCFAs (Fig. 3e), were reduced in uveitis patients compared to healthy controls. This supports a role for SCFAs in regulation of human uveitis.

### 4. Akkermansia negatively correlates with Firmicutes in both sEAU and clinical uveitis

Among all taxa in HuFl mice, Akkermansia abundance was shown to be a prominent correlate of severe uveitis and IFN-γ-producing CD4+ T cells in the gut (Fig. 2e-f, Extended Data Fig. 3a–b), whereas SCFA-producing Firmicutes were reduced in the same animals (Fig. 2e, g). We therefore analyzed their relative abundance and potential interdependence to determine whether competitive interactions between these taxa might contribute to autoimmune uveitis.

We first quantified abundance of Akkermansia and Firmicutes, and found them to be inversely correlated (Fig. 4a). An in-depth computational analysis of the bacterial association network confirmed the negative correlation between Akkermansia and multiple families of SCFA-producing Firmicutes (Fig. 4b, Extended Data Fig. 3c). Levels of major SCFAs in feces also exhibited an inverse correlation with Akkermansia abundance (Fig. 4c). Taken together, this supports an interpretation of a negative interaction between Akkermansia and a range of SCFA-producing microbes.

**Fig. 4.**
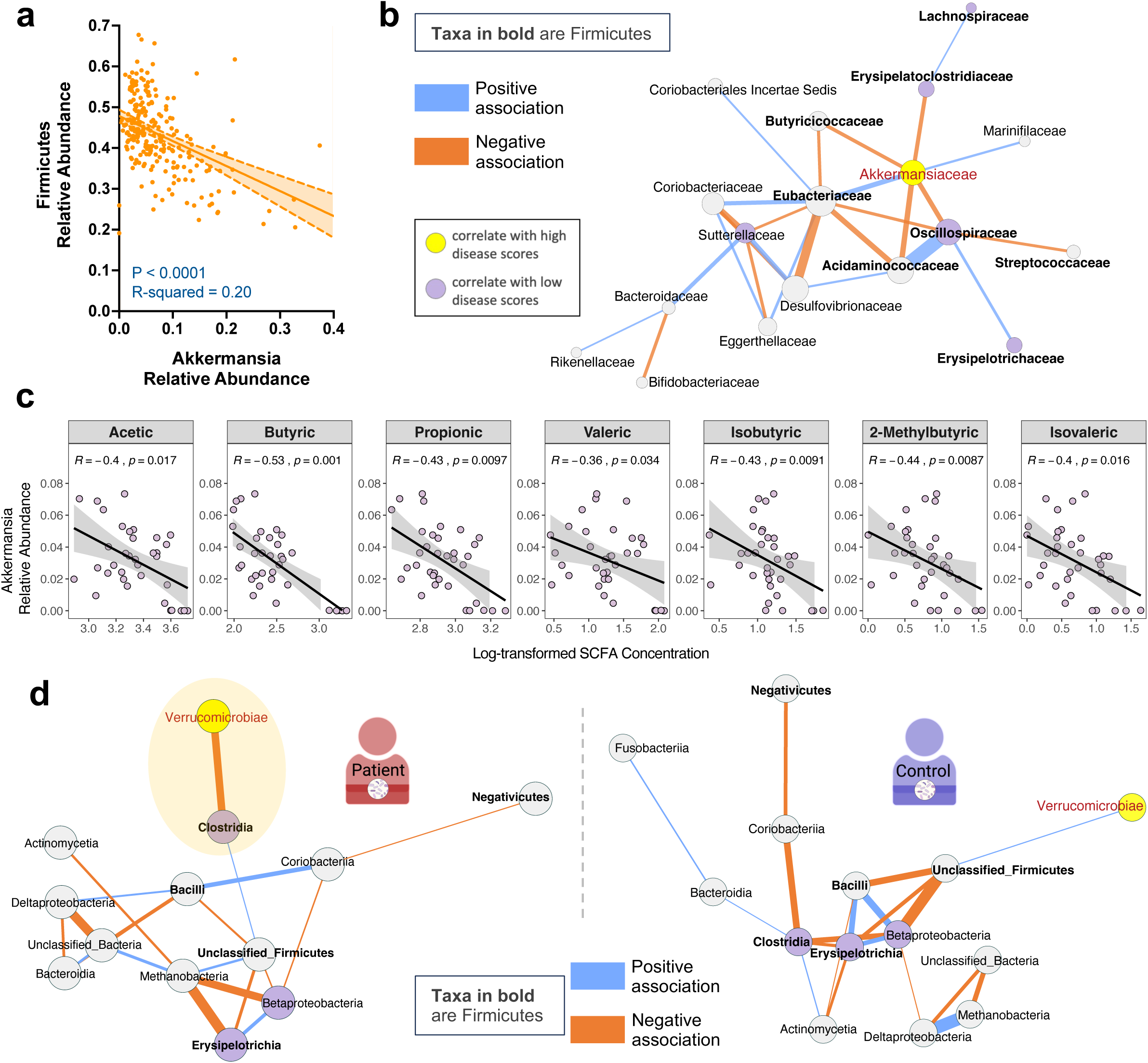
Akkermansia negatively correlates with Firmicutes in both sEAU and clinical uveitis. **a**, Simple linear regression showing a negative correlation of microbial relative abundance between Akkermansia (equivalent to Verrucomicrobia in this dataset) and Firmicutes in HuFl R161H mice (*n =* 234, UVW combined, weanling and adult samples included). **b**, Fecal bacterial association network exemplified in HuFl cohort V (*n =* 79, weanling and adult samples included). Network constructed at the taxonomic level of the bacterial family. Correlations with a threshold of Pearson r > 0.5 were transformed into dissimilarities via the “unsigned” distance metric, on which edge weights (strength of correlations) are based. Blue edges correspond to positive associations and orange edges to negative ones. Yellow and purple nodes represent taxa enriched in high *vs.* low disease categories (FDR < 0.05), with Firmicutes labeled as bolded taxon names. Bigger nodes are more influential in the network (higher eigenvector centrality). **c**, Spearman correlation (visualized with a linear trend line) showing a negative association between relative Akkermansia abundance and SCFA concentrations in R161H gut (*n =* 35, UVW and SPF combined). Akkermansia was selected following multivariable association between SCFA concentrations and microbial features using MaAsLin2. **d**, Class level association networks in treatment-naïve uveitis patients (left, *n =* 8) and healthy controls (right, *n =* 12) from the same cohort of autoimmune uveitis. Correlations with a threshold of Pearson r > 0.3 are shown. Node and edge color schemes same as in **b**. Shaded ellipse in the patient network highlights the negative correlation between Verrucomicrobiae (Akkermansia) and Clostridia (Firmicutes), and this relationship is absent from the healthy human gut.

Similarly, gut microbiome interaction networks of uveitis patients displayed a negative relationship between Verrucomicrobia (primarily Akkermansia) and SCFA-producing Firmicutes (Clostridia, class level), but this relationship was not evident in healthy subjects (Fig. 4d). Together with the functional and metabolite findings in Fig. 3d–e, this compositional relationship supports the relevance of the sEAU model to clinical disease.

### 5. Akkermansia enrichment increases uveitis and reduces Firmicutes relative abundance and circulating SCFAs

To validate the proposed microbiome interaction network *in vivo*, we used a gain-of-function approach, where GF R161H mice received either an inoculum of Akkermansia or Akkermansia mixed with HuFl mouse donor fecal flora. In the first paradigm, we associated GF mice *de novo* with cultured *Akkermansia muciniphila* (the type species of Akkermansia) or with sterile PBS as control, and maintained them in static microisolators using sterile handling techniques (Extended Data Fig. 4a). This paradigm was designed to generate an Akkermansia-enriched, low-complexity community rather than a strict mono-association.

Consistent with this design, mice developed gut communities of very low diversity (≤10 detectable taxa per mouse), dominated by Verrucomicrobia (Akkermansia) and Firmicutes (Extended Data Fig. 4b-d).

First paradigm: A single oral gavage led to successful and sustained engraftment of Akkermansia, and resulted in higher uveitis scores (Extended Data Fig. 4e-f). Akkermansia was nearly undetectable in the PBS control group, whose gut microbiota became dominated by Firmicutes (Extended Data Fig. 4g). As such, the relative abundances of Akkermansia and Firmicutes showed a near-perfect inverse relationship (r-squared > 0.99, p < 0.0001) (Extended Data Fig. 4h). Other bacterial phyla that are prominent in normal mouse or human gut flora were low or on the borderline of detectability (Extended Data Fig. 4i). Thus, this model reflects a minimal-diversity gut ecosystem dominated by the two taxa that we hypothesize to display competitive dynamics that control expression of disease.

Second paradigm: To examine this hypothesis under more physiological conditions, where Akkermansia is not given a major “head start”, we associated GF R161H mice with flora from HuFl mice (selected for low disease and low abundance of Akkermansia) “spiked”, or not, with cultured *A. muciniphila* (Fig. 5a). Fecal microbiome profiling confirmed successful microbial engraftment and stability of Akkermansia-high or Akk-low status (Fig. 5d & Extended Data Fig. 5a). Akkermansia “spiked” flora induced more severe uveitis than base HuFl alone (Fig. 5b-c) and the relative abundances of Akkermansia and Firmicutes were consistently inversely correlated (Fig. 5e). Notably, among the major bacterial phyla, Firmicutes was the only phylum whose abundance was decreased in Akk-enriched mice (Fig. 5d & Extended Data Fig. 5b), particularly those taxa considered “professional” SCFA producers^24^ (Extended Data Fig. 5c). To confirm that the effects described above were indeed Akkermansia-specific, we associated GF R161H mice with the same HuFl “spiked” with *E. coli*. The disease-promoting phenotype and the reduction in Firmicutes abundance were not recapitulated, supporting the notion that the observed effects were specific to Akkermansia (Extended Data Fig. 5d-f). Together, these results reinforce a dynamic balance between Akkermansia and Firmicutes that modulates autoimmune uveitis in a physiologically relevant setting.

**Fig. 5.**
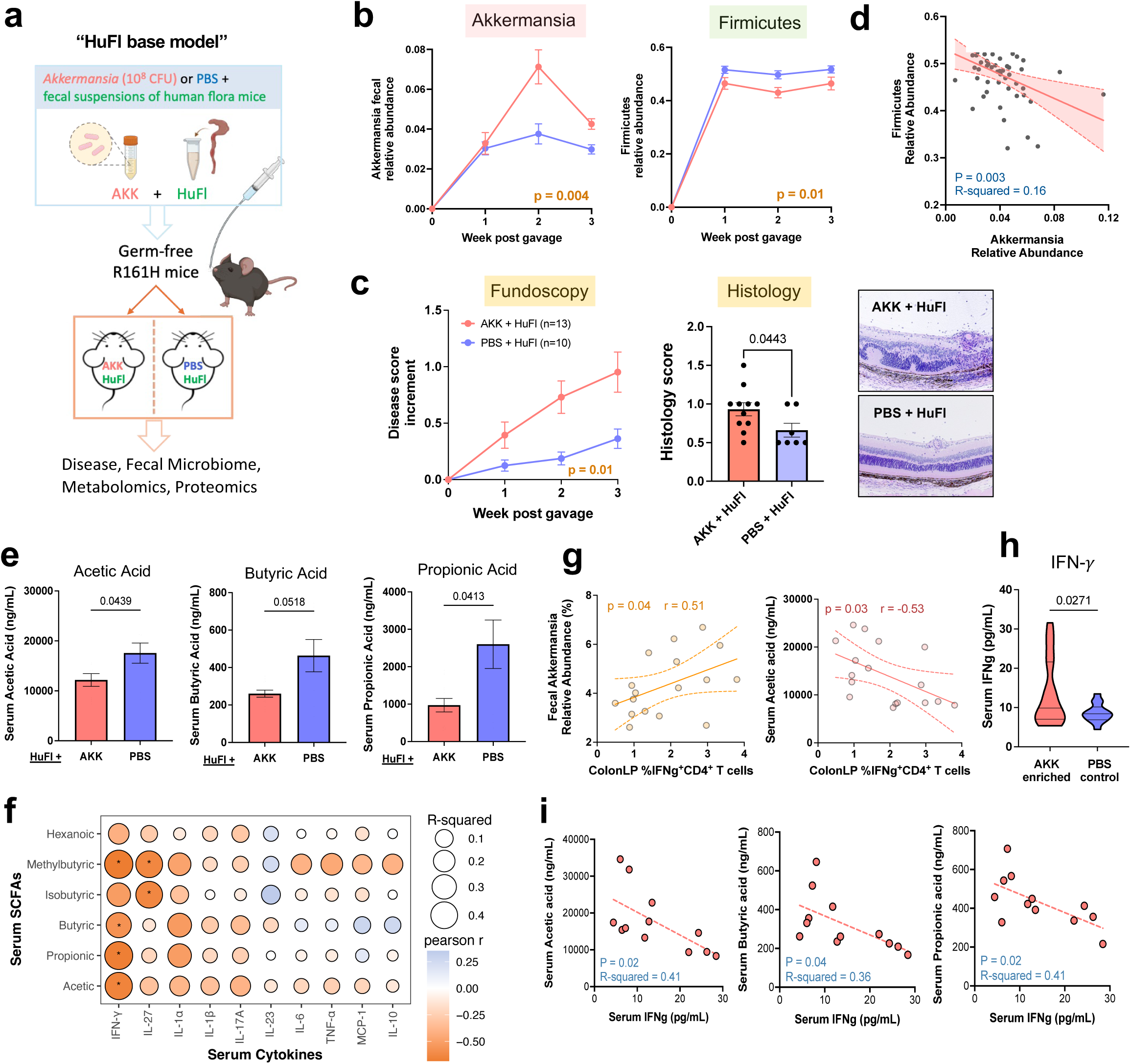
Akkermansia promotes uveitis by outcompeting SCFA producers and decreasing circulating SCFA levels. **a**, Study design of microbe reconstitution experiments. Previously GF R161H mice gavaged with a baseline human gut microbiota suspension (HuFl) plus *A. muciniphila* culture (AKK, 10^8^ CFU) or sterile PBS. **b**, Relative abundance of Akkermansia and Firmicutes in feces from one representative of 3 experiments (*n =* 7 and 6 for Akk-rich and control, respectively). **c**, Left: Fundoscopy score increments from starting disease scores, combined from three experiments. Right: Histology scores and example images from 2 out of 3 experiments (*n =* 11 and 7 for Akk-rich and control, respectively). Mean values ± SEM. P-value determined by Welch-corrected t test. **d**, A negative correlation of relative abundance between Akkermansia and Firmicutes (*n =* 52, AKK and PBS groups combined, multiple time points included) by simple linear regression. **e**, Serum concentrations of main SCFAs are significantly lower in Akk-rich (*n =* 13) mice than in PBS controls (*n =* 10). Serum harvested 3 – 4 weeks post gavage, three experiments combined. **f**, Correlational heatmap of SCFA levels with cytokine concentrations in serum of engrafted mice. *P < 0.05, modeled by simple linear regression as well as Pearson correlation. **g**, Spearman correlation (visualized with a linear trend line) showing Akkermansia relative abundance associated positively, and serum acetic acid associated negatively, with the frequency of IFN-*γ*-producing CD4 T cells in colonic LP. **h**, Akk-enriched (*n =* 18) mice had higher serum IFN-*γ* concentrations compared to PBS control mice (*n =* 14), five experiments combined, of which two experiments used the minimal flora model. **i**, Representative trend lines show that main SCFA concentrations are negatively associated with IFN-*γ* levels in serum. Data in **g, i** are from 2 experiments (*n =* 12) using the minimal flora model.

To mechanistically link Akkermansia–Firmicutes dynamics with circulating SCFA levels available to regulate uveitis-relevant immune responses, we measured serum SCFAs and host immune parameters. All three major SCFAs – acetic, butyric and propionic acids – were reduced in Akk-high mice compared with Akk-low mice (Fig. 5f), consistent with the negative relationship between Akkermansia and Firmicutes. This reduction was not observed in the *E.coli* control group (Extended Data Fig. 5g). Given the known immunomodulatory role of SCFAs^21^, we examined whether this metabolomic shift was associated with an altered systemic effector response. In both the Akk-associated minimal flora model and the Akk-enriched HuFl model, reduced serum SCFAs were tightly coupled with elevated serum and/or gut IFN-γ responses (Fig. 5g–j). These findings suggest that reduced SCFA availability may contribute to the enhanced Th1 response associated with Akkermansia enrichment.

The final mechanistic question was whether Akkermansia can provide surrogate antigenic stimuli to trigger activation of the R161H TCR-expressing cells in the gut^2^. To address this, we used an *in vitro* activation assay, where naïve R161H CD4 T cells and antigen-presenting cells were cultured with Akk-rich HuFl gut content extracts, or with heat-killed Akkermansia. Unlike SPF gut flora extracts or the cognate peptide antigen^2,10^, neither Akkermansia preparation activated the retina-specific T cells, as measured by expression of CD69 and Nur77 (Fig. 6a-b), indicating that Akkermansia was not acting as an antigen mimic under these assay conditions.

**Fig. 6.**
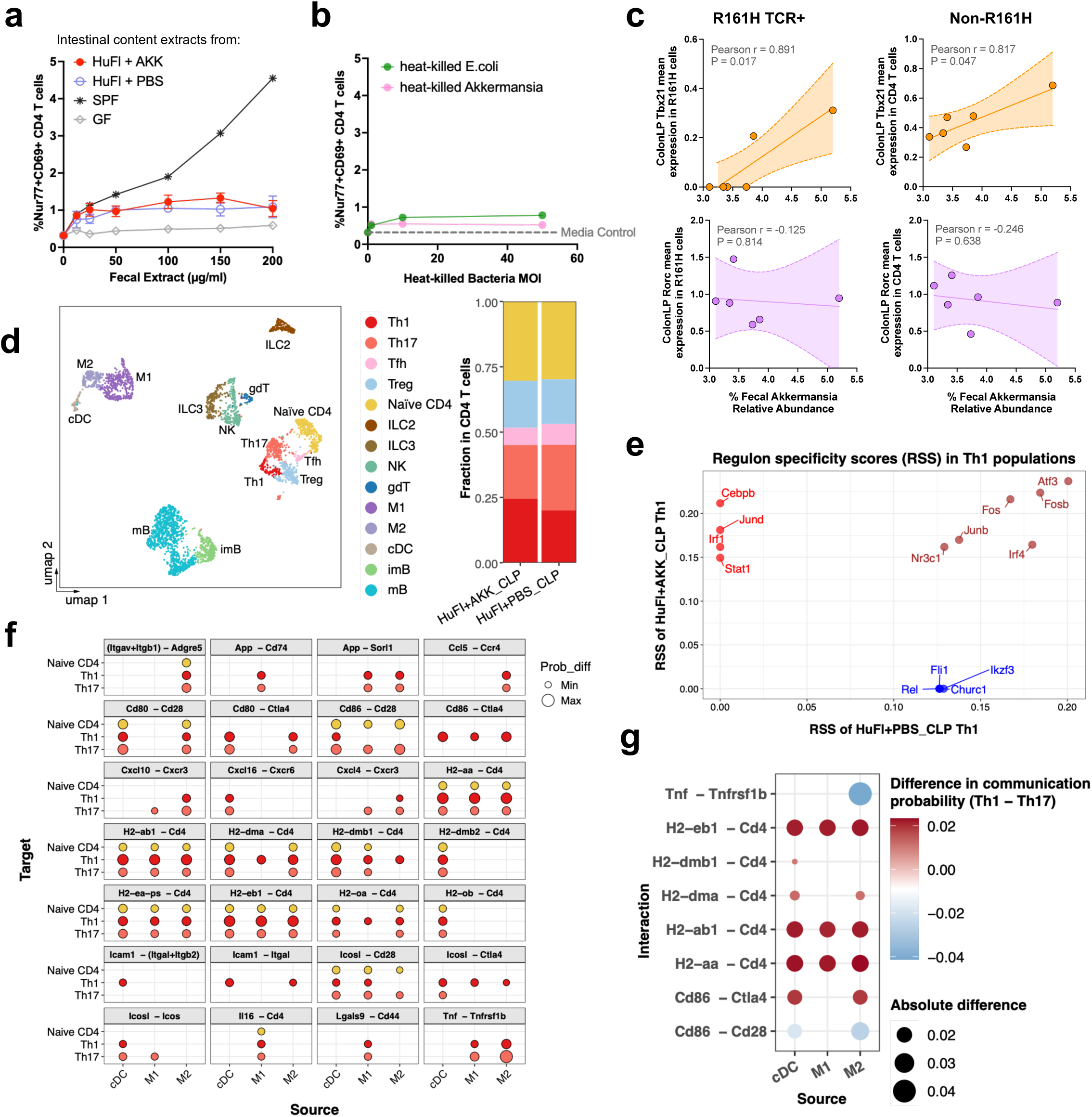
Akkermansia does not directly stimulate IRBP-specific T cells but can upregulate Th1 responses in the Akk-rich gut. **a**–**b**, Induction of Nur77 and CD69 expression in naïve IRBP-specific CD4 T cells from *Tcra^-/-^*R161H-*Nr4a1^GFP^* mice after 22 hours of stimulation with splenic dendritic cells (DC) from WT mice. Intestinal content extracts from R161H mice of indicated microbiome groups (**a**), and heat-killed Akkermansia or *E.coli* (**b**) were used as stimulants. GF (grey line in **a**) and media only (dotted line in **b**) represent baseline levels of activation. Multiplicity of Infection (MOI) is relative to DC. *n =* 4, 3 for HuFl+AKK and HuFl+PBS groups, respectively. **c**, Correlation of Akkermansia relative abundance at week 3 post-gavage with the normalized transcript abundance (*Tbx21* or *Rorc*) in R161H TCR-expressing and non-R161H CD4 T cells in the colon LP (CLP). Data from 5’ single cell RNAseq from the Akkermansia reconstitution experiment, combined from HuFl+AKK (*n =* 3) and HuFl+PBS (*n =* 3) groups. **d,** UMAP of CD45^+^ cells from CLP of a representative mouse from each group, and proportions of CD4^+^ T cell subpopulations. Tfh, T follicular helper cells; ILC2/3, innate lymphoid cells; NK, natural killer cells; gdT, gamma delta T cells; M1/M2, macrophages; cDC, conventional dendritic cells; imB/mB, immature/mature B cells. **e,** Top 10 regulons enriched in Th1 cells in the CLP from representative HuFl+PBS (blue and brown) *vs.* HuFl+Akk (red and brown). Brown indicates regulons shared between groups. **f,** CellChat-inferred cell-cell communication in the CLP showing increased signaling from cDC and macrophages to naïve CD4 T and effector Th1 and Th17 cells in HuFl+Akk compared to HuFl+PBS. **g**, Differential signaling increases in Th1 cells compared to Th17 cells in HuFl+Akk gut.

Single cell RNA-seq analysis of colonic CD4 T cells, of which an average of 15% expressed the R161H TCR, revealed that both R161H TCR-expressing and non-R161H (endogenous) T cells had a Th1, rather than a Th17, signature, that positively correlated with abundance of Akkermansia (Fig. 6c). Akk-high colonic CD4 T cells showed an increased proportion of Th1 cells with enriched interferon-and inflammation-associated transcriptional programs (Stat1, Irf1, Cebpb, Jund), indicating an augmented Th1 transcriptional profile (Fig. 6d-e). CellChat inferred broadly increased myeloid-to-T-cell communication (Fig. 6f), including greater predicted MHC class II-associated APC–CD4 interaction scores involving Type 1 than Type 17 cells (Fig. 6g).

VDJ analysis did not reveal evidence for a 2^nd^ TCR among R161H TCR+ cells, and *in silico* analysis did not reveal shared sequence(s) between Akkermansia and IRBP161-180, which is the cognate R161H TCR epitope, arguing against direct R161H TCR activation by Akkermansia under the conditions tested. Together, these findings support a model in which Akkermansia regulates disease and shapes the T-cell effector response at 2 levels: potentially through innate, adjuvant-like mechanisms modulating APC-T cell interactions, as well as indirectly, by modulating SCFA-producing Firmicutes. This working model is depicted in Figure 7a.

**Fig. 7.**
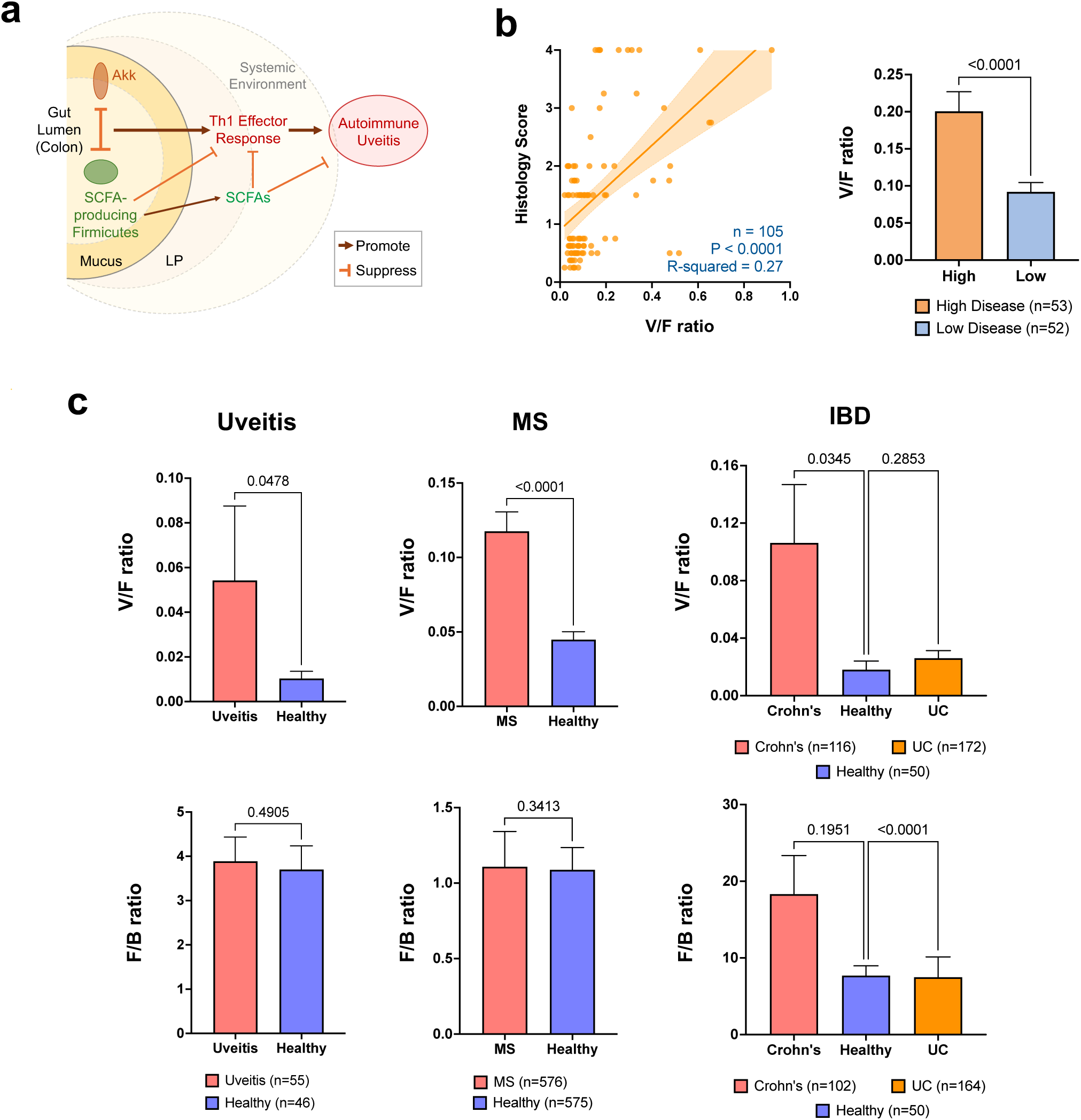
The inverse relationship between Verrucomicrobia and Firmicutes extends to clinical uveitis, multiple sclerosis (MS) and Crohn’s disease. a, Graphical summary of the working model. b, A positive correlation between V/F (Verrucomicrobia/Firmicutes) ratios and histology scores of HuFl mice by simple linear regression, and V/F ratios compared between high and low disease of HuFl mice. c, V/F and F/B (Firmicutes/ Bacteroidetes) ratios compared between patients from uveitis, MS and IBD and their respective healthy controls. Analyzed datasets of MS and IBD (Crohn’s disease and UC) and healthy controls were from public sources. Uveitis and MS cohorts included those established on treatment. For F/B ratio in the IBD cohort, samples with <0.1% relative abundance of either Bacteroidetes or Firmicutes were removed from the analysis. Data are presented as mean values ± SEM. Statistical significance was determined by Mann-Whitney U test (one-tailed analysis was applied for uveitis and MS) and Kruskal–Wallis test with P-values adjusted for post hoc multiple comparisons (for IBD).

### 6. The inverse relationship between Verrucomicrobia and Firmicutes extends to clinical uveitis, multiple sclerosis and Crohn’s disease

Because effector Th1 skewing in the Akk-rich gut environment was observed in both R161H and endogenous CD4 T cells, we examined whether a related microbial pattern occurred in human inflammatory conditions, such as uveitis, multiple sclerosis (MS) and inflammatory bowel disease (IBD). To establish a common metric for comparing healthy to diseased state, we used the phylum-level ratio of Verrucomicrobia / Firmicutes (V/F ratio) which consistently tracked with disease severity in HuFl mice (Fig. 7b). Application of this metric to the entire cohort of our clinical uveitis samples (so as to minimize any selection bias), revealed higher V/F ratios in uveitis patients compared to healthy subjects (Fig. 7c). We next examined publicly available MS^25^ and IBD^26^ datasets, the latter comprising patients with Crohn’s disease and ulcerative colitis (UC). Compared to healthy controls, the V/F ratio in MS, and in Crohn’s disease patients––a condition predominantly driven by Th1-mediated inflammation––but not in UC patients^27,28^, was markedly elevated (Fig. 7c). Taken together, the gut microbial interaction networks indicated a negative correlation of Verrucomicrobia (Akkermansia) with Firmicutes in R161H mice and uveitis patients; V/F comparisons showed a related pattern in MS and Crohn’s disease datasets.

## DISCUSSION

In this study, we report a gut microbial interaction network that regulates autoimmune and inflammatory processes along the “gut–eye axis”. By integrating data-driven analyses and *in vivo* microbial perturbation, we provide compelling evidence for a negative relationship between Akkermansia and Firmicutes and link it to disease severity. Specifically, Akkermansia appears to promote sEAU in HuFl R161H mice by outcompeting SCFA-producing Firmicutes in the gut, resulting in diminished SCFA levels and enhanced Th1 responses both locally and systemically (Fig. 7a). Our findings provide a mechanistic basis for how the gut microbiome impacts ocular autoimmunity, dissect the complex interacting stimuli that underpin this process, and highlight its broadly generalizable nature.

Our network-based perspective contrasts with previous studies, which have mostly focused on how individual gut microbial taxa influence disease and host immunity^4^. *A. muciniphila* has been implicated across a spectrum of inflammatory and autoimmune diseases^29^. However, different reports have described context-dependent and sometimes divergent functional effects. For example, Akkermansia has been reported to promote Th1 polarization of human PBMCs^30^, and its relative abundance correlated with disease activity in MS patients^25,30–32^. In contrast, in vivo Th1-polarizing activity has not been observed in Akk-monocolonized mice^30^ and T cell responses to Akkermansia in SPF mice have not correlated with a consistent T cell effector phenotype^33^. At the same time, *A. muciniphila* has been associated with disease attenuation in specific models, including delayed diabetes development in NOD mice^34^ and reduced DSS-induced colitis and colitis-associated tumorigenesis following administration of pasteurized *A. muciniphila* or its outer-membrane protein Amuc_1100^35^. These observations, together with our finding that Akkermansia affects sEAU through interactions that modulate the broader microbial network, support the notion that the immunological impact of Akkermansia—and of individual microbes more generally —depends strongly on microbial community context and host factors.

The R161H sEAU model provides a platform to dissect these contextual effects. When the model was first established, we identified signaling by a (microbiota-dependent) mimic antigen in the SPF flora of R161H mice as an important, but possibly not the only, disease trigger^2^. In the case of Akkermansia as a disease-promoting influence that does not appear to act as a surrogate antigen, it is likely that other HuFl member(s) may provide this function. The preferential enhancement of MHC-II–associated APC–CD4 interactions in Akkermansia-enriched gut is conceptually reminiscent of the SFB-driven Th17 program^36,37^, in which dendritic-cell MHC-II presentation and commensal-specific TCR selection create a Th17-polarizing milieu that instructs adaptive effector differentiation. Akk seems to promote a polarizing milieu encouraging Th1 programming of CD4+ cells without evidence for IRBP-specific microbial antigen mimicry.

We did not attempt to identify the HuFl microbial mimic, as identifying it would lack translational relevance. The R161 TCR recognizes a mouse class II (I-A^r^)-restricted epitope of IRBP. Human MHC class II molecules would almost certainly present a different epitope(s) from any given antigen than does mouse I-A^r^, plus the epitope(s) would differ between individuals, given the heterogeneity of human MHC haplotypes. Moreover, whereas IRBP is a dominant retinal autoantigen in mice, human uveitis more commonly involves recall responses to arrestin^1^. Therefore, although the HuFl R161H sEAU model cannot reveal specific antigenic targets involved in human uveitis, it can reveal conserved mechanisms shared across species, such as the microbial interaction network identified in this study.

A key feature of this microbial network is the negative relationship between Akkermansia and Firmicutes, which links gut microbial composition to systemic inflammation. This antagonism likely arises from multiple, non-mutually exclusive mechanisms. Both taxa are prominent colonizers of the colon and may engage in direct niche competition and indirect metabolic interactions. Akkermansia promotes mucus degradation during gut inflammation^38,39^. The compromised mucus barrier can facilitate microbial encroachment and amplify mucosal inflammation^40^, while the resulting inflammatory environment can increase epithelial oxygenation and oxidative stress^41^, creating inhospitable conditions for obligate anaerobes like Clostridia^42^. Finally, while Akkermansia enrichment led to a reduction in SCFA-producing Firmicutes in our system, SCFAs can in turn act to suppress Akkermansia growth and mucolytic activity^43^. These reciprocal interactions provide a plausible basis for the context-dependent effects of Akkermansia reported across disease models and support our interpretation that its impact on autoimmunity is determined not by the organism in isolation, but by the microbial and host ecosystem in which it resides.

All those interactions are reflected in the V/F ratio, which may serve as an indicator of inflammatory state and immune polarization across species, supporting its translational value. A similar “legacy” metric Firmicutes to Bacteroidetes (F/B ratio) has been used to indicate overall shifts in gut microbiome composition, but is now being phased out due to its inconsistent disease associations^44,45^. We propose that the V/F ratio is more specifically informative in autoimmune and inflammatory settings.

Of note, numerous studies report an association between uveitic diseases and gut inflammation. Uveitis is associated with an increased risk of Crohn’s (but not of UC)^46^, and Crohn’s patients are more likely than UC patients to develop uveitis^47^. The findings are compatible with the notion that the V/F ratio increases under conditions where gut inflammation co-occurs with extraintestinal autoimmunity. The V/F ratio may also rise under Th1-dominant inflammatory conditions, as observed in our data. Our analyses reveal elevated V/F ratios not only in our patients with uveitis, but also in published datasets from patients with MS^25^ and IBD^26^. Within IBD, increases in V/F were evident in Crohn’s disease (Th1-dominant^48–50^), but not in UC (Th2-like^51–53^).

Limitations of the study: The complexity of the Akkermansia–Firmicutes axis poses a major challenge to resolving microbial relationships at species and strain levels. Emerging approaches such as bacterial spatial transcriptomics^54^ are not yet sufficiently developed to dissect mechanisms by which Akkermansia antagonizes SCFA-producing Firmicutes. That said, outcompetition could involve physical elimination via bacteriocins acting on SCFA producers^55^. In addition, the asynchronous nature of disease development in individual mice and the low breeding rate of GF R161H mice constrains the scale and breadth of experiments that can feasibly be performed.

In conclusion, our findings show that a complex microbial interaction network—rather than individual taxa—plays a central role in shaping the gut–eye axis. The Akkermansia– Firmicutes relationship likely represents a generalizable ecological principle in which microbial competition influences systemic immune balance and susceptibility to autoimmunity and inflammation, and which extends across species. Future studies aimed at defining the mechanisms by which Akkermansia gains an advantage over SCFA-producing Firmicutes, may guide the development of microbiome-based interventions for autoimmune and inflammatory diseases that intersect with gut dysregulation.

## METHODS

### Mice

R161H mice on the B10.RIII background^10^ were bred in-house and maintained under SPF conditions and fed standard facility chow *ad libitum*. GF and HuFl R161H mice were maintained in the gnotobiotic isolators and fed CMF diet (Oriental Yeast Co., Ltd., Japan), irradiated 50kGy with Cobalt-60. For the sterile and static housing of gnotobiotic mice in the SPF facility (described ahead), the same CMF diet and autoclaved water were provided. The animal study protocol (NEI-688) was approved by the NEI Institutional Animal Care and Use Committee.

### Generation of gnotobiotic mice with human stool samples

Human stool samples of healthy volunteers were collected at Keio University according to the study protocol^56^ approved by the Institutional Review Boards (approval number 20150075) and informed consent was obtained from each subject. Stool samples were suspended in 20% glycerol in PBS, snap frozen in liquid nitrogen and stored at-80°C. The U, V, and W stocks were thawed and gavaged into GF R161H mice and individual cohort lines were established in separate gnotobiotic isolators at Sankyo Labo Service Corporation, Inc. (Ibaraki, Japan). The V and W cohorts were successfully generated by gavage and breeding. Because mice gavaged with the U stock failed to produce viable offspring, new GF mice were introduced into the isolator and co-housed to generate the U cohort. After colonies were established, HuFl mice were transported from Japan to NEI in the germ-free shipping containers.

### Evaluation of uveitis phenotype

Development of uveitis was monitored by fundoscopic observation via a binocular microscope of anesthetized mice at the indicated age. For histopathology, eyes were enucleated and fixed in 4% glutaraldehyde for 1 h and transferred to 10% formaldehyde for additional 24 h, embedded in methacrylate, and processed for H&E staining. Disease scores of fundoscopy and histology were assigned by a masked observer on a scale of 0–4, according to the criteria for EAU scoring described in detail elsewhere which are based on the number, size and type of lesions^57^.

### HuFl R161H fecal suspension preparation

Cecal and colonic contents from HuFl mice of varying disease severity were collected under sterile conditions and immediately transferred on ice to an anaerobic chamber (Coy Laboratory Products). Contents were filtered three times through a 70 µm cell strainer in 20% glycerol–PBS, and aliquoted to prevent repeated freeze–thaw cycles. Each sample was prepared from either individual mice or pools of mice with the same disease severity. The baseline HuFl suspension for reconstitution experiments was prepared from two mice with low uveitis scores and validated by 16S sequencing to contain ∼1% Akkermansia (ranges 1–8% in tested samples).

### In vivo microbial enrichment by gnotobiotic reconstitution

*A. muciniphila* or *E. coli* were cultured in mucin-BHI medium inside the anaerobic chamber or in LB medium in the aerobic incubator, respectively. Microbial monocultures were prepared in advance, and the first passage was timed so that cultures reached exponential growth phase immediately prior to oral gavage. On the day of gavage, GF R161H were transferred from the gnotobiotic isolators to sterile and static microisolator cages with Reemay filter tops.

Cages were opened only within a biosafety cabinet and maintained using aseptic handling procedures. For fundoscopic examination, mice were briefly removed from the cabinet for <1 min, then promptly returned to their cages. Mice were assigned to experimental or control groups based on initial uveitis scores (<1), age (4–7 weeks), and sex. While mice were recovering from anesthesia, gavage material was prepared inside the anaerobic chamber for *A. muciniphila*, in the aerobic bacterial hood for *E. coli*, or as sterile PBS, which in the “HuFl base model” was combined with the selected HuFl fecal suspension, so each mouse in the experimental group would receive an additional approximately 10^8^ – 10^9^ CFU of the microbe of interest in 200uL HuFl fecal suspension via oral gavage. For each gavage experiment, *A. muciniphila* cultures were initiated in advance so that they reached the empirically determined exponential-growth range at the time of harvest (OD600 = 1.1–1.3 under our culture conditions). Cells were harvested immediately before preparation of the gavage inoculum. Because a reliable OD-to-CFU conversion was established only for *E. coli*, the CFU of *A. muciniphila* administered was determined retrospectively by plating. Specifically, a homogeneous aliquot of the *A. muciniphila* suspension used to prepare the gavage inoculum was retained on ice inside the anaerobic chamber. After gavage was completed, this aliquot was serially diluted and plated, and colonies were enumerated after 4–5 days of anaerobic culture to determine the CFU administered in each experiment. Similarly, *E. coli* was harvested during the empirically determined exponential-growth phase, typically a few hours after the initial subculture, at an OD600 of 0.6–1.0. CFU/mL was estimated from OD600 using the Agilent *E. coli* Cell Culture Concentration from OD600 Calculator, and the inoculum was adjusted to the desired dose before gavage. Fecal sampling and fundoscopy scoring were performed on a weekly basis under sterile protocols.

### Fecal DNA extraction and quantification of microbes

Fecal pellets were collected from individual mice, immediately frozen on dry ice and stored at-80°C until bacterial DNA extraction in house as previously described^58^ or by the NCI Genetics and Microbiome Core. Absolute abundance of bacteria in each fecal sample was assessed by quantitative polymerase chain reaction (qPCR) using Akkermansia or *E. coli* standard curves with Fast SYBR Green Supermix (Applied Biosystems, Waltham, MA, USA) and run on a QuantStudio^TM^ 7 Pro Real-Time PCR System (Applied Biosystems). Specific primers for Akkermansia were Akk.mu3.Fwd (5’-GCGTAGGCTGTTTCGTAAGTCGTGTGTGAAAG-3’) and Akk.mu3.Rvs (5’-GAGTGTTCCCGATATCTACGCATTTCA-3’)^59^, and for *E. coli* were Uni515F (5’-GTGCCAGCMGCCGCGGTAA-3’) and Ent826R (5’-GCCTCAAGGGCACAACCTCCAAG-3’)^60^. Five ng of fecal DNA was amplified per 10 µL reaction with the following cycling conditions: initial denaturation at 95 °C for 20 sec, followed by 40 cycles of 95 °C for 3 sec and 63 °C for 30 sec. A melt-curve analysis was included at the end of each run to confirm amplification specificity. Bacterial counts (CFU) per 5 ng of DNA were extrapolated from fitted standard curves correlating Cq values with bacterial quantity.

### 16S rRNA gene sequencing and analyses

Fecal samples were processed at the NCI Genetics and Microbiome Core for 16S rRNA gene sequencing as detailed in prior work^58^. 16S amplicon sequence variants (ASV) were inferred with YAMS16 v1.7 (https://github.com/jhbadger/YAMS16), a pipeline running DADA2^61^, and taxonomically classified using the SILVA reference database (release 138.1)^62^. Microbial diversity, differential abundance and multivariable association analyses were performed with phyloseq v1.42.0^63^, MaAsLin2 v1.12.0^64^, microViz v0.12.3^65^ in R. Microbial association networks were analyzed with NetCoMi^66^.

### Shotgun metagenomic sequencing and analyses

Selected DNA samples were submitted to CosmosID, Inc. (Germantown, MD, USA) for shallow shotgun sequencing (3 million reads per sample). Specifically, DNA libraries were prepared using the Nextera XT DNA Library Preparation Kit (Illumina, Inc., San Diego, CA, USA) and IDT Unique Dual Indexes with total DNA input of 1 ng. Genomic DNA was fragmented using a proportional amount of Illumina Nextera XT fragmentation enzyme. Unique dual indexes were added to each sample followed by 12 cycles of PCR to construct libraries. DNA libraries were purified using AMpure magnetic Beads (Beckman Coulter, Inc., Brea, CA, USA) and eluted in QIAGEN EB buffer. DNA libraries were quantified using Qubit 4 fluorometer and Qubit™ dsDNA HS Assay Kit. Libraries were then sequenced on the Element AVITI platform 2×150 bp. JAMS v1.9.9^67^ and HUMAnN v3.9^68^ were used for taxonomic and functional profiling, and genes involved in metabolic pathways of the main SCFAs (acetate, butyrate and propionate) were identified based on InterPro, Gene Ontology and Pfam annotations, and those present in the uveitis datasets were compared between diseased and healthy groups.

### Targeted metabolomics and cytokine quantification

Cecal and colonic contents (collectively referred to as “fecal contents”), eyes, and serum were collected from age-matched mice at harvest. Eyes were fixed for histological analysis as described above. Fecal and serum samples were snap-frozen on dry ice and stored at −80 °C. For metabolite profiling, at least 100 mg of fecal contents and 150 µL of serum per mouse were transferred to designated tubes and shipped on dry ice to Metabolon, Inc. (Morrisville, NC, USA) for absolute quantification of eight short-chain fatty acids (acetic, propionic, butyric, isobutyric, 2-methylbutyric, valeric, isovaleric, and hexanoic acids).

Serum cytokine concentrations were measured using the Mouse Inflammation Panel (13-plex, LEGENDplex™, BioLegend, San Diego, CA, USA) according to the manufacturer’s instructions. Data were acquired on a CytoFlex LX flow cytometer (Beckman coulter) and analyzed with LEGENDplex Data Analysis Software (BioLegend) to quantify cytokine levels based on standard curves.

### Tissue harvesting and flow cytometry analyses

Peripheral lymphoid tissues (submandibular lymph nodes, spleen, peripheral lymph nodes, mesenteric lymph nodes), and gut lamina propria from small intestine and colon were collected and prepared for single cell suspensions as described elsewhere^2^ from R161H mice aged between 7 and 11 wk-old. For the intracellular markers, cells were stimulated ex vivo with 50 ng/ml PMA and 1 µg/ml Ionomycin (MilliporeSigma) in the presence of BD GolgiPlug protein transport inhibitor (containing Brefeldin A) for 4 hours, washed, fixed with 4% paraformaldehyde (Electron Microscopy Sciences) in PBS and permeabilized with BD Perm/Wash buffer prior to the staining. Cells were stained with fluorochrome-conjugated antibodies (clones) from BD Biosciences, BioLegend or eBioscience/Thermo Fisher Scientific. Based on our previous titration experiments, all the antibodies were used at the dilution of 1:1000. Surface staining panel: FITC-TCR beta chain (H57-597), BB700-CD69 (H1.2F3), PE-CD62L (MEL-14), PE/Dazzle594-CD183/CXCR3 (CXCR3-173), PE/Cy7-CD199/CCR9 (CW-1.2), Alexa Fluor 700-CD8a (53-6.7), APC-eFluor 780-CD44 (IM7), Brilliant Violet (BV) 421-CD103 (2E7), BV510-CD45 (30-F11), BV650-CD4 (RM4-5), BV785-CD196/CCR6 (29-2L17), BUV496-CD3 (145-2C11). Intracellular staining panel: Alexa Fluor 488-IL-17F (9D3.1C8), PerCP/Cyanine5.5-T-bet (4B10), PE-RORgt (Q31-378) PE/Dazzle594-IL-10 (JES5-16E3), PE/Cy7-GM-CSF (MP1-22E9), APC-IL-22 (IL22JOP), Alexa Fluor 700-Foxp3 (FJK-16s), APC-eFluor 780-CD8a (53-6.7), BV421-IFN-g (XMG1.2), BV510-CD4 (RM4-5), BV650-IL-17A (TC11-18H10), BV786-CD44 (IM7), BUV496-CD3 (145-2C11). Dead cells were excluded by staining with ViaKrome 808 Fixable Viability Dye from Beckman Coulter. Fluorescence-activated cell sorting analyses were done using CytoFlex LX (Beckman coulter). FlowJo v10 (BD Biosciences) was used for data analyses.

### In vitro activation assay on retina-specific T cells

IRBP-specific T cells were harvested from spleen and peripheral lymph nodes of *Tcra^-^*^/-^ *Nr4a1^EGFP^* (Nur77^GFP^) R161H mice and were enriched for naïve CD4^+^ cells by negative selection using Mouse Streptavidin Rapidsphere Isolation Kit (EasySep, StemCell Technologies). The following biotin-conjugated anti-mouse antibodies (clones) from BioLegend, BD Biosciences were used at 1:250 dilutions: CD8 (53-6.7), CD11b (M1/70), CD11c (N418), CD16/32 (93), CD19 (6D5), CD24 (M1/69), CD25 (PC61), CD49b (DX5) CD69 (H1.2F3), CD45R/B220 (RA3-6B2), Ly-6G/Ly-6C (Gr1, RB-8C5), I-A/I-E (M5/114.15.2), NK1.1 (PK136), *γ*δTCR (GL3), Ter 119 (TER-119). CD11c^+^ dendritic cells were purified from spleen of WT B10.RIII mice as antigen-presenting cells using CD11c MicroBeads Ultrapure (Miltenyi Biotec). T cells and dendritic cells were co-cultured at a 4:1 ratio in the presence of heat-killed *A. muciniphila* (95°C for 5 min), heat-killed *E. coli* (60°C for 30 min), or bacteria-rich extracts from the gut content^2^. T cell activation was assessed after 22 h by flow cytometric analysis of CD69 and Nur77 (GFP) expression.

### 5’ single cell RNAseq sample processing

Cells from colon LP were labeled with TotalSeq C anti-mouse HashTags 1-10 (C0301-C0310, clones M1/42; 30-F11) and 14 cell surface antibodies from BioLegend at the dilution of 1:100. Cell surface anti-mouse antibodies: CD4 (DNA_ID C0001, clone RM4-5), CD8a (C0002, 53-6.7), Ly-6C (C0013, HK1.4), CD11b (C0014, M1/70), Ly-6G (C0015, 1A8), CD90.2 (C0075, 30-H12), CD19 (C0093, 6D5), CD11c (C0106, N418), F4/80 (C0114, BM8), I-A/I-E (C0117, M5/114.15.2), NK-1.1 (C0118, PK136), TCRβ chain (C0120, H57-597), TCR g/d (C0121, GL3), and CD3 (C0182, 17A2). Labeled cells were sorted for live CD45^+^ and CD4^+^ populations with PE anti-mouse CD45.2 (clone 104, BioLegend), eFluor^TM^ 450 anti-mouse CD4 (clone RM4-5, eBioscience) and 7-AAD (eBioscience) using FACSAria III/Fusion sorters (BD Biosciences). Post-sort viability exceeded 80%, and hash-tagged samples were pooled at equal ratios within each experimental group, combining CD45^+^ and CD4^+^ fractions at 1:1 ratio to enrich for CD4^+^ T cells. Pooled cells were resuspended in PBS with 0.04% BSA and loaded onto a 10x Genomics Chromium X instrument at 700–1200 cells/µL for single cell 5’ gene expression and TCR profiling. Libraries were prepared using the Chromium Next GEM Single Cell 5’ Reagent Kits v2 (Dual Index) following the manufacturer’s protocol, and sequenced on NovaSeq 6000 platform (Illumina, Inc., San Diego, CA, USA).

### Single cell RNAseq data analyses

Six mice (three from each group) were used for analyses. Sequencing data were processed with Cell Ranger v9.0.1 (10x Genomics) for demultiplexing and alignment to the Mus musculus reference (GRCm39, Genome Reference Consortium, 2024). Cell doublets were detected and removed using Scrublet v0.2.3^69^. Cells exhibiting high mitochondrial transcript content (≥ 5%) or abnormal gene counts (<500 or> 8000) were excluded. Expression data were log-normalized and scaled using the Scanpy v1.11.0^70^ in Python v3.10. Dimensionality reduction was performed using Principal Component Analysis (PCA) followed by Uniform Manifold Approximation and Projection (UMAP), and cell clusters were identified with the Leiden algorithm^71^. Cell type annotation was based on computational classification using SingleR v2.8.0^72^ with mouse immune reference data from celldex::ImmGenData v1.16.0 in R v4.4, as well as the expression of canonical cell type marker genes, and validated by cell lineage–specific surface antibody. The expression of both Trav16 and Trbv5 in a CD4 T cell was the standard used to identify IRBP-specific T cells^10^. Seurat v5.1.0^73^ was also used to generate object files for downstream application of CellChat^74^ and SCENIC^75^ in the R environment. CellChat was applied to infer changes in the strongest cell-cell interactions between Th1 cells and other cell types in the colon LP in Akk-rich mouse. SCENIC was used to reconstruct gene regulatory networks and to assess transcription factor activity at the single-cell level, enabling identification of Akkermansia-associated alterations in Th1 cell programming and lineage maintenance.

### NEI clinical samples and public human data

Stool samples were obtained from patients with autoimmune uveitis and healthy controls enrolled in a clinical cohort at the NEI, NIH. All participants provided written informed consent under the protocol approved by the Institutional Review Board (Protocol 13-EI-0072). A stratified cohort of treatment-naïve autoimmune uveitis, with no systemic therapy during the preceding two years, was selected for in-depth gut microbial network analysis, representing primarily posterior-and pan-uveitis cases and ocular-restricted phenotypes such as idiopathic uveitis and birdshot chorioretinopathy. 16S rRNA gene and shotgun metagenomic sequencing of human fecal samples were performed by the Microbiome Cores at the NCI and the Goodman-Luskin Microbiome Center (Los Angeles, CA, USA). Public microbiome data of IBD patients’ stool are from PRJNA1198911^26^, and that of MS patient stool from the International MS Microbiome Study (iMSMS)^25^.

### Statistical analyses

Statistical analyses were conducted with R v4.4, Python v3.10, and GraphPad Prism v10.5.0. Non-parametric tests such as Mann-Whitney U and Kruskal-Wallis tests were used for most comparisons between two or more groups. Group differences in multivariate community structure were evaluated by Permutational Multivariate Analysis of Variance (PERMANOVA). Pearson’s or Spearman’s rank correlation, as well as multivariable association (MaAsLin2 and equivalent methods) were used to assess associations between microbial features and disease or immune phenotypes. P-values were adjusted for multiple comparisons using the false discovery rate (FDR) where applicable, and statistical significance was indicated as follows: * P < 0.05, ** P < 0.01, *** P < 0.001, and **** P < 0.0001. Data in bar plots are presented as mean values ± SEMs.

### Data availability

Relevant sequencing data have been deposited under BioProject ID PRJNA1451962.

### Code availability

No custom code was developed for this study. All analyses were performed using publicly available software packages, and the specific versions and parameters used are described in the Methods section.

## Author contributions

A.Z., R.H. and R.R.C. conceived and designed the study. R.R.C and R.H. supervised the project. A.Z., R.H., Y.J., A.G., S.A., and C.E.M. performed experiments and collected data. A.Z., J.H.B., Z.W., G.S., and V.N. performed computational analyses. S.H., H.N.S., J.W.L. and K.H. provided human microbiome samples, and S.K. and H.N.S. contributed to clinical interpretation. X.X., M.J.M., Z.P., B.X. provided experimental input and technical assistance. R.P. and N.M. provided technical expertise in bacteriology. J.H.B., Z.W., J.A.M., J.P.J. and C.O. contributed analytical tools, resources and expertise. A.Z. and R.H. analyzed and interpreted data and prepared figures. A.Z., R.H. and R.R.C. wrote and finalized the manuscript. All authors discussed the results, reviewed, and approved the final manuscript.

## Acknowledgments

We are grateful to NEI and NHLBI Flow Cytometry Cores, NEI Histology and Genetic Engineering Cores, Ms. Wuxing Yuan, Dr. Shah Rashed and members of NCI Microbiome Core, UCLA Goodman-Luskin Microbiome Center Microbiome Core, and all members of the Caspi Lab for their invaluable experimental assistance. We extend special thanks to Dr. Katsuko Sudo (Tokyo Medical University, Japan), Mr. Hirofumi Kobayashi and staff at Sankyo Labo Service Corporation, Inc. (Japan), as well as the gnotobiotic management team from NEI Veterinary Research and Resources Section for their expert support and guidance in GF and gnotobiotic colony maintenance. We also thank Dr. Eduard Ansaldo (NIAID) for sharing protocols for Akkermansia culture, Dr. Kikuji Itoh (Japan SLC, Inc.), Dr. Koji Atarashi and members of the Honda Lab for providing experimental advice. Finally, we acknowledge Dr. Geoffrey A. Mueller and staff at NIEHS Nuclear Magnetic Resonance Group for their generous assistance with metabolomics, and Drs. Kelsey Wheeler, Daniel Sellers, and Reuben Allen (Massachusetts Institute of Technology) for their insightful discussions and critical feedback. This study was supported by NIH/NEI Intramural funding, Project Number EY000184.

## Commercial Disclosures

None

