## Supplemental Figures for "Gut microbial interaction networks control autoimmunity to neuroretina"

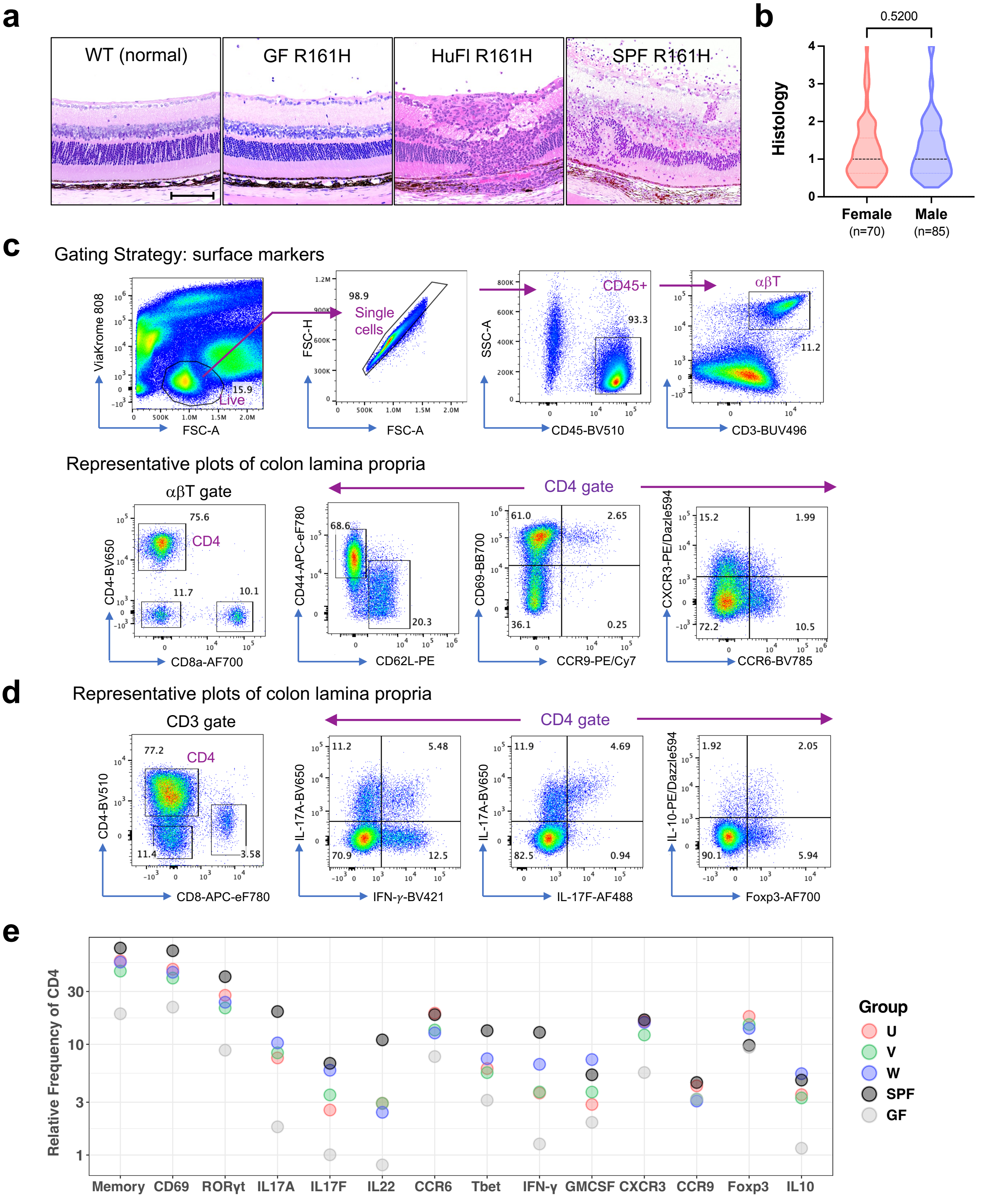


**Extended Data Fig. 1 | Disease and CD4 T cell phenotypes of HuFl R161H. a**, Representative retinal histology images contrasted between WT (normal retina) and R161H (GF, HuFl-V and SPF). The scale bar in the WT image shows 100 µm. **b**, Histology scores are comparable between female and male HuFl R161H mice at ages of 7-11 weeks old. Data combined from UVW cohorts (Mann-Whitney U test, $p = 0.52$). **c**, Gating strategy and representative flow cytometry plots of the surface markers from colon LP. **d**, Representative flow cytometry plots of the intracellular markers from colon LP. **e**, Mean relative frequency of each cell surface or intracellular marker of colon LP CD4 T cells, differentially colored by the microbiome groups.

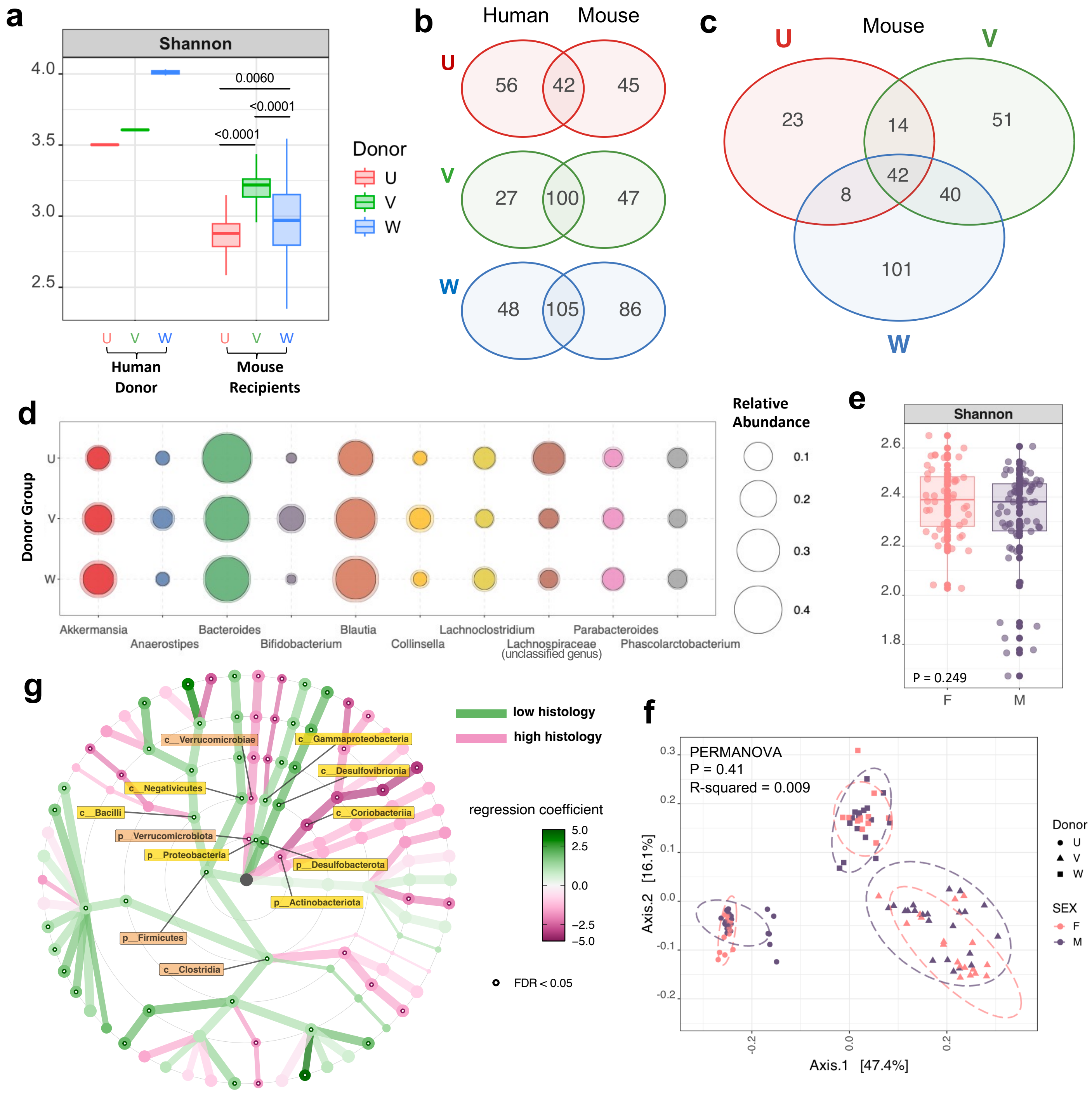


**Extended Data Fig. 2 | Characterization of HuFl R161H gut microbiota and its disease association. a**, Alpha diversity (Shannon Index) of gut content samples from mouse recipients, juxtaposed with stool samples from human donors. P < 0.0001 from Kruskal–Wallis test between mouse recipient groups, with p-values from Dunn's multiple comparisons shown in the plot. *n* = 77, 79, 78 for U, V, W mice, respectively. **b–c**, Venn Diagrams showing numbers of shared and unique bacterial taxa in fecal microbiome between each human and the recipient mouse cohort (**b**), and among three mouse cohorts (**c**). **d**, Average bacterial relative abundance in three mouse cohorts. Top 10 most abundant genera are shown. **e,** Alpha diversity (Shannon Index) of adult microbiome between female and male HuFl R161H (Wilcoxon rank sum test, p = 0.249). **f**, Beta diversity of adult HuFl R161H microbiome based on Bray–Curtis dissimilarity shows clustering by donor cohort rather than by sex. **g**, Multivariate taxonomic association with general linear models visualized as a dendrogram, showing bacterial taxa enriched in high (*n* = 16) or low (*n* = 24) histology categories in adult V-cohort R161H. Differentially abundant phyla and classes are labeled, FDR < 0.05. For **a–b**, each human donor sample has two technical replicates, and for **b–d**, *n* = 50, 59, 64 for mouse cohort U, V, W, respectively (original gavaged recipients and several offspring generations combined). For **e–f**, *n* = 44 and 61 for female and male, respectively (UVW combined).

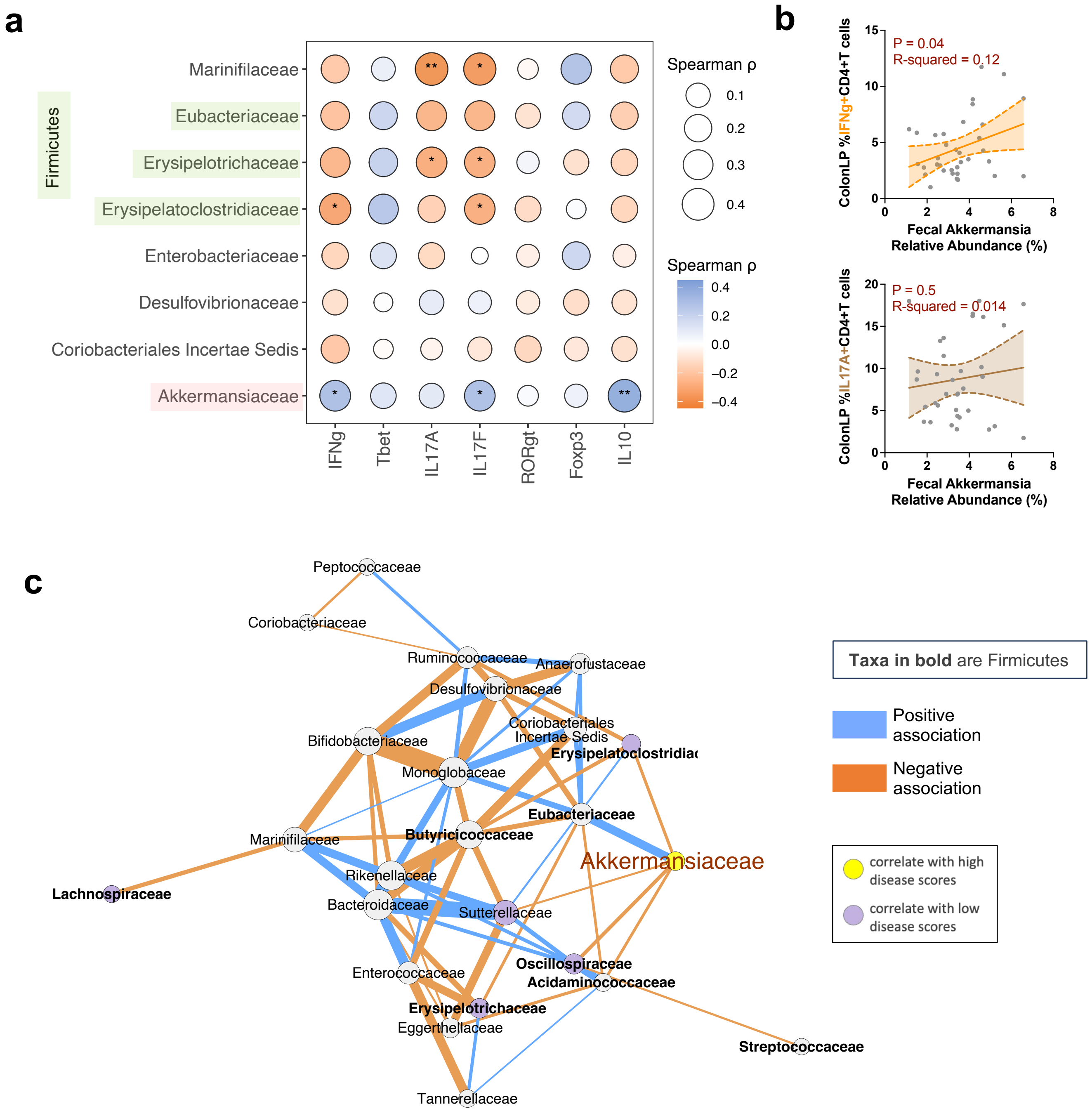


**Extended Data Fig. 3 | Akkermansia – Firmicutes axis in HuFl R161H mice. a**, Correlational heatmap showing associations of most abundant bacterial families (Firmicutes in green shade) with relative frequencies of colon LP CD4 T cells expressing each intracellular marker in adult HuFl R161H ($n$ = 99, UVW combined). *P < 0.05, **P < 0.01, modeled by Spearman's rank correlation with Spearman's ρ defining the color gradient and circle size. **b**, Representative trend lines with 95% confidence interval (shaded area) show that relative Akkermansia abundance was positively associated with IFN-$\gamma$-producing, but not with IL-17A-producing, CD4 T cells in colon LP. **c**, Bacterial (family level) association networks in HuFl R161H mice ($n$ = 234, UVW combined). Correlations with a threshold of Pearson r > 0.35 are shown. Node and edge color schemes same as in **Fig. 4** (node colors indicate statistically significant disease associations, FDR < 0.05).

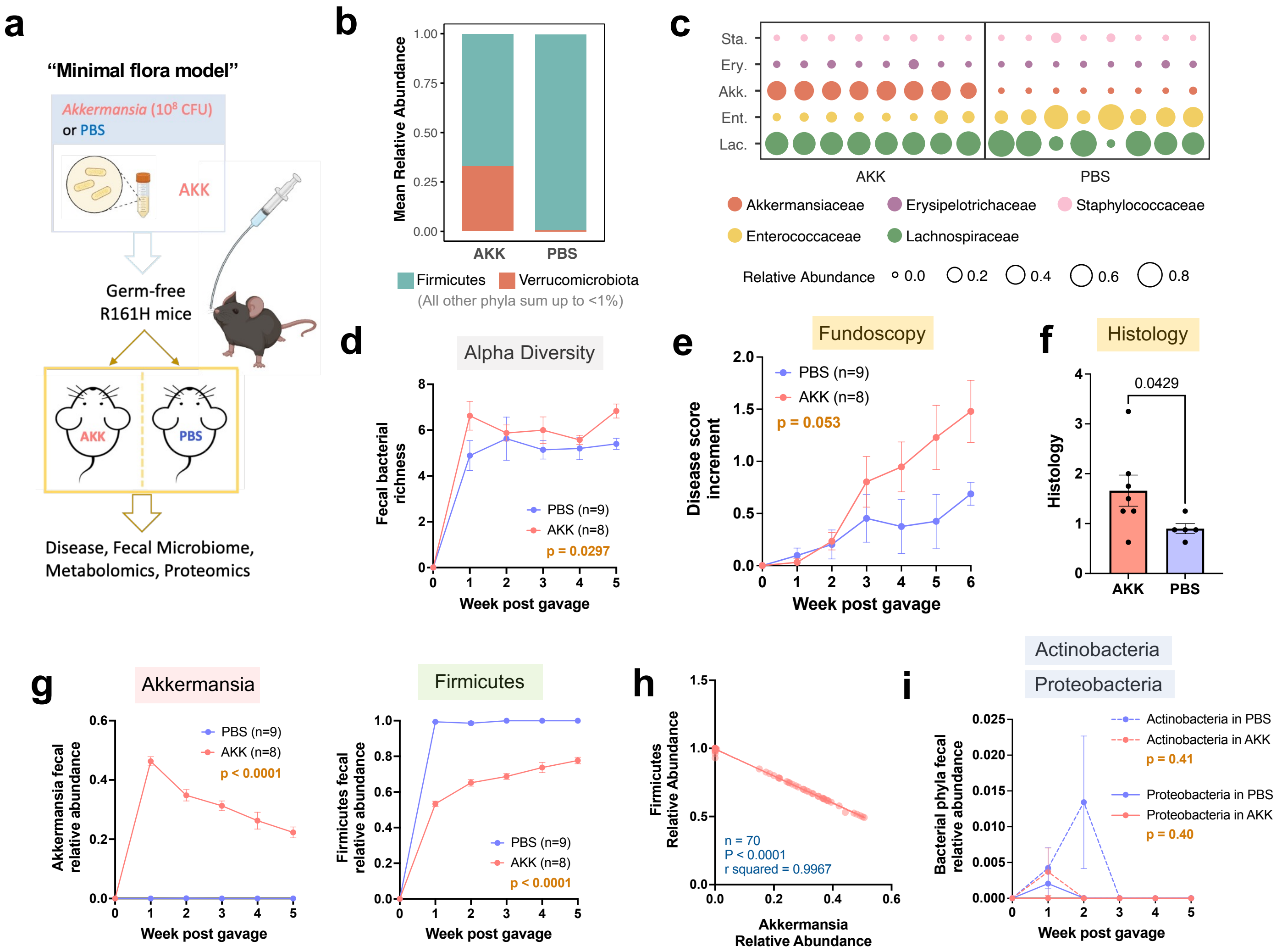


**Extended Data Fig. 4 | Akkermansia promotes uveitis in the context of a minimal gut flora.**
**a**, Experimental design of the "minimal flora model". **b**, Mean relative abundance of the two dominant fecal bacterial phyla, Firmicutes and Verrucomicrobiota, in AKK ($n$ = 8) and PBS ($n$ = 9) groups. **c**, Relative abundance of the top five fecal bacterial families in individual mice (represented by each column), two weeks after reconstitution. **d**, Fecal bacterial richness, a measure of alpha diversity, compared between AKK and PBS groups. **e**–**f**, Fundoscopy score increments from starting disease scores (**e**) and histology scores (**f**) of AKK and PBS mice. **g**, Relative abundance of Akkermansia and Firmicutes in feces, compared between AKK and PBS groups. **h**, A negative correlation of relative abundance between Akkermansia and Firmicutes by simple linear regression ($n$ = 70, AKK and PBS groups combined, multiple time points included). **i**, Relative abundance of Actinobacteria and Proteobacteria in reconstituted mice. Bacteroidetes, another bacterial phylum that is normally abundant in the mouse gut, was nearly undetectable in these mice. Data are presented as mean values ± SEMs. P-value determined by Mann-Whitney U test in (**f**) and two-way ANOVA in (**d**, **e**, **g**, **i**). Results combined from two experiments. In plots with time series, microbial abundance at the time point "0" (GF status) represents a theoretical zero.

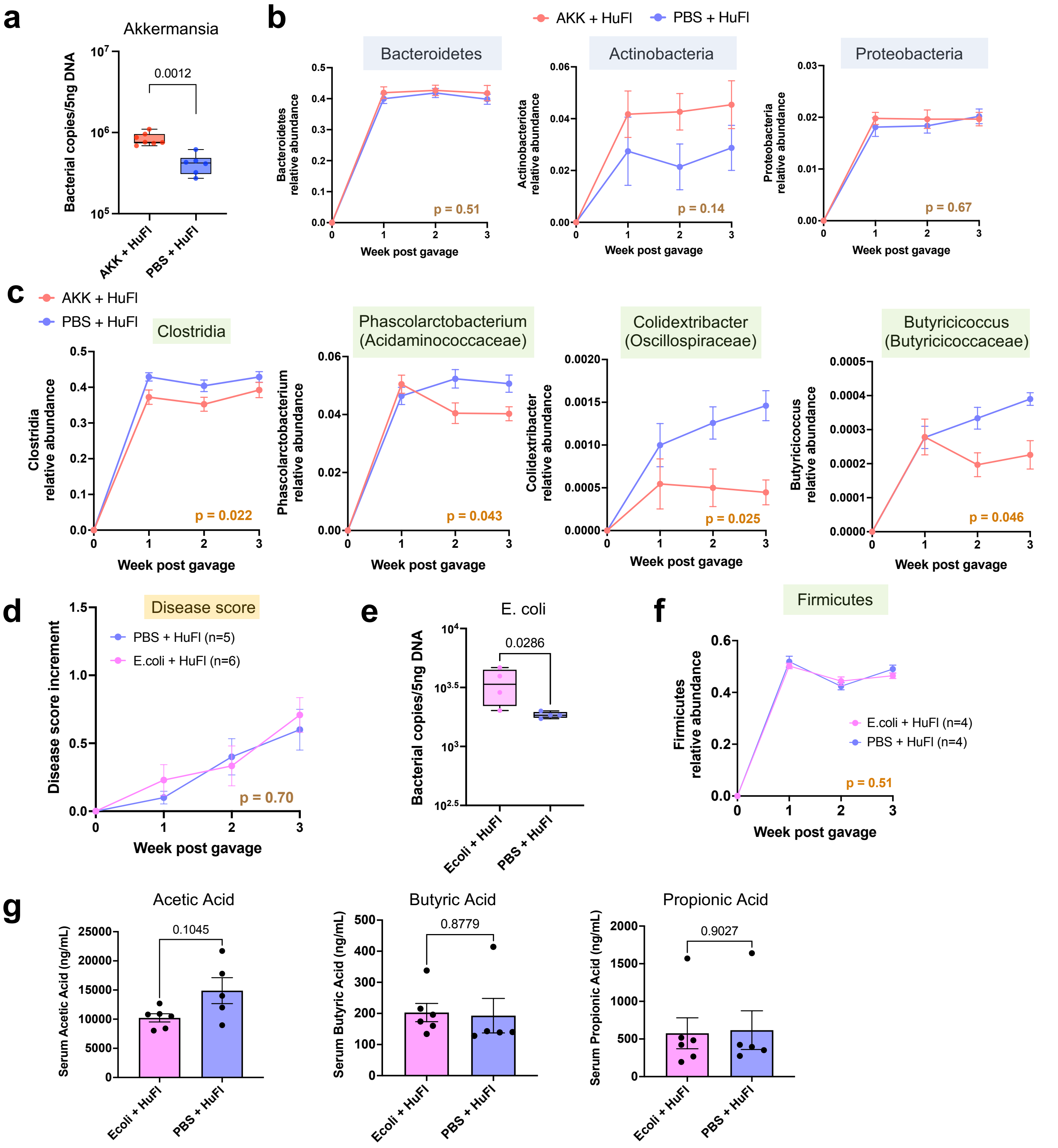


**Extended Data Fig. 5 | Akkermansia promotes uveitis, while *E. coli* does not, in concert with base human flora.** **a**, Absolute abundance of Akkermansia compared between Akk-rich and control mice two weeks after reconstitution. **b**, Relative abundance of major phyla Bacteroidetes, Actinobacteria and Proteobacteria **c**, Relative abundance of representative SCFA-producing Firmicutes (class to genus levels) in reconstituted mice (one representative experiment, $n$ = 7 and 6 for Akk-rich and control, respectively). Data are presented as mean values ± SEMs. Statistical comparison between experimental groups was conducted using two-way ANOVA and p-values are shown in each graph. **d–g**, HuFl + E.coli ($10^8$ CFU) reconstitution experiments, as controls to Akkermansia. **d**, E.coli group did not develop more severe uveitis than PBS controls. **e**, Absolute abundance of E.coli one week after reconstitution. **f**, Relative abundance of Firmicutes. **g**, Serum concentrations of main SCFAs in E.coli-rich and control mice. Serum harvested four weeks post gavage. Data are presented as mean values ± SEM. Statistical comparison between experimental groups was performed using two-way ANOVA or Welch's t test.
